# A Taxonomy-Informed Sparse DNA Foundation Model for Microbial Genomics

**DOI:** 10.64898/2026.09.22.753215

**Authors:** Athan Z. Li, Shiyuan Wang, Shupeng Cheng, Yuxuan Du, Ruishan Liu

## Abstract

Microorganisms are crucial to Earth’s ecosystems, with genomes encoding functions important to agriculture, biotechnology and human health. Although genomic language models have advanced DNA representation learning, the extensive diversity of microorganisms and highly imbalanced taxonomic composition of existing pretraining corpora pose challenges for effective microbial genomic sequence modeling. Here we present MicroGlot, a taxonomy-informed microbial DNA foundation model pretrained on 3.70 million sequences comprising 378.3 billion nucleotides from 99,700 species. MicroGlot represents hierarchical relationships among taxa using hyperbolic embeddings and integrates this knowledge into a sparse mixture-of-experts architecture. Probing frozen embeddings across layers showed that MicroGlot captures both microbial phenotypic traits and taxonomic identity. Compared with a taxonomy-ablated variant pretrained under the same scheme, incorporating taxonomic knowledge consistently improved representation quality across model depth. Combining sparse computation with efficient architectural and training techniques, MicroGlot achieved competitive probing and fine-tuning performance with low computational requirements. Species-level expert-routing fingerprints aligned more closely with taxonomic groups than tetranucleotide-frequency profiles, indicating that expert usage recapitulates taxonomic structure. These results show that combining sparse architecture with taxonomic knowledge enables efficient and biologically informative genomic language modeling across diverse microbial taxa. MicroGlot is publicly available at https://huggingface.co/athanzli/MicroGlot.

## 1 Introduction

Microorganisms underpin biological processes and applications spanning plant health^1^, sustainable agriculture ^2^, industrial and environmental biotechnology^3,4^, and human health and disease ^5^. Characterizing these roles often requires identifying the microorganisms involved, for which genomic sequences provide critical references^6,7^. Yet extracting biologically meaningful information from DNA sequence remains challenging ^8^. Genomic language models (gLMs) ^9^ have emerged as a general framework for addressing this problem by learning transferable representations from large collections of genomic sequences through self-supervised pretraining. Early models such as DNABERT adapted bidirectional transformer pretraining to DNA and learned contextual representations for regulatory sequence prediction ^10^. HyenaDNA introduced an alternative convolution-based architecture that extended sequence modeling to contexts of up to one million nucleotides^11^. Like DNABERT^10^, it was pretrained only on the human reference genome. Pretraining coverage has since expanded across increasingly diverse genomic sources. DNABERT-2 ^12^, DNABERT-S^13^, the Nucleotide Transformer (NT) ^14^, GENA-LM ^15^, and OmniNA ^16^ used multispecies pretraining, while ProkBERT^17^ and Evo ^8^ focused on microbial genomic sequences. Evo2 further extended genome-scale pretraining across all domains of life ^18^.

Expanding pretraining across species requires gLMs to accommodate substantial phylogenetic diversity. The scale of this diversity is reflected in genome-based resources such as the Genome Taxonomy Database (GTDB), which organizes 901,341 bacterial and archaeal genomes into 199,923 species clusters ^19,20^. At the same time, taxonomic groups can be highly unevenly represented in gLM pretraining corpora. For example, among the bacterial, fungal, and protozoan genomes in the NT multispecies corpus, bacterial sequences contribute substantially more nucleotides than fungi and protozoa combined^14^. Learning from sequences that are both phylogenetically diverse and unequally represented therefore places competing demands on a shared set of model parameters. An analogous challenge arises in multilingual language modeling, where increasing language coverage in a dense transformer of fixed capacity can lead to competition for shared capacity and interference between languages, a phenomenon termed the curse of multilinguality ^21,22^. This trade-off is further influenced by data imbalance, as increased sampling of high-resource languages can improve their performance at the expense of lower-resource languages ^21^. Sparse mixture-of-experts (MoE) models ^23^ are an effective approach to alleviating these capacity constraints by activating only a subset of model parameters for each token, thereby increasing model capacity without a proportional increase in per-token computation. In multilingual pretraining, Switch Transformers improved performance over dense mT5 baselines across 101 languages ^23^, while sparsely gated MoE models in No Language Left Behind (NLLB) improved the trade-off between cross-lingual transfer and interference, particularly for low-resource languages ^24^. These findings motivate the use of sparse MoE architectures for learning from taxonomically diverse and disproportionately represented microbial genomic sequences.

Alongside architectural approaches for accommodating microbial genomic diversity, annotated taxonomy provides structured biological knowledge that can guide learning across taxa. RefSeq^25^, for example, links reference sequences to taxonomic annotations from NCBI Taxonomy^26^, while the International Committee on Taxonomy of Viruses (ICTV) provides standardized taxonomic assignments for viruses ^27^. Beyond species identity alone, these resources encode hierarchical relationships among organisms that reflect their evolutionary organization. Previous genomic models have begun to incorporate species information during sequence learning. SpeciesLM^28^ and SPACE^29^ use learned species embeddings to condition DNA sequence modeling, while Evo^8^ and Evo2 ^18^ incorporate taxonomic lineage information through sequence conditioning tokens. However, learned species embeddings treat species identities as trainable representations without constraining their geometry to reflect the hierarchical relationships present in a taxonomic tree. Explicitly encoding these relationships may therefore provide a complementary source of biological structure for multispecies genomic representation learning across diverse microbial taxa.

Here we introduce MicroGlot, a taxonomy-informed gLM that integrates hierarchical taxonomic representations with a sparse MoE architecture. We assembled a pretraining corpus of 3.70 million sequences comprising 378.3 billion nucleotides from 99,700 species spanning bacteria, archaea, fungi, protists, and viruses, together with plasmid sequences. From the resulting taxonomy, we derived Poincaré embeddings ^30^ that represent hierarchical relationships among taxa in a hyperbolic space, providing taxonomic context for sequence representation learning and expert routing. To further improve computational efficiency, the backbone incorporates modern language-model components such as grouped-query attention (GQA) ^31^ and FlashAttention-2^32^.

We evaluated whether MicroGlot learns biologically informative representations by training supervised probes on frozen embeddings from each model layer to predict microbial taxonomic identity and phenotypic traits, including growth and environmental preferences. MicroGlot achieved strong probing performance while using the smallest pretraining token budget among the evaluated gLMs. An ablation study further showed that incorporating taxonomic information consistently improved representation quality across model depth relative to an otherwise matched model pretrained without taxonomic knowledge. We next assessed computational efficiency through inference benchmarks across increasing sequence lengths and downstream fine-tuning on microbial taxonomic classification and epigenetic marks prediction tasks. MicroGlot combined competitive predictive performance with low inference and fine-tuning costs, demonstrating the computational advantages of its optimized architecture. Finally, analysis of expert routing revealed taxonomically structured expert usage, with related microbial taxa exhibiting similar routing patterns. Together, these results show that integrating hierarchical taxonomic knowledge with sparse genomic language modeling enables efficient and biologically informative representation learning in microbial genomics.

## 2 Results

### 2.1 A diverse microbial genomic sequence corpus for taxonomy-informed pretraining

To represent the extensive microbial diversity ^19^, we sought sequences spanning major cellular microbial domains and viral realms with broad species coverage, while requiring annotations across multiple taxonomic ranks. We therefore combined curated reference sequences from RefSeq^25^ (release 232), virus isolates and their classifications from the ICTV Virus Metadata Resource^27^ (MSL40 v1) and plasmid sequences with host annotations from PLSDB^33^ (version 2024_05_31_v2).

Lineages follow NCBI Taxonomy^26^ for RefSeq sequences and PLSDB hosts and the ICTV classification for virus isolates, spanning domain or realm, kingdom, phylum, class, order, family, genus and species (Methods). After sequence quality filtering and sampling to limit species overrepresentation (Methods), the corpus encompassed 99,700 species, with broad taxonomic structure reflected in sequence composition (Fig. 1a). These species were represented by 3,695,065 sequences and 378.3 billion nucleotides (Fig. 1b). Its taxonomic coverage spanned the three cellular microbial domains Bacteria, Archaea and Eukaryota and seven viral realms, including Adnaviria, Duplodnaviria, Monodnaviria, Riboviria, Ribozyviria, Singelaviria and Varidnaviria. It covered 67,476 bacterial, 1,299 archaeal, 13,084 fungal, 898 protist and 16,692 virus species, together with plasmids assigned to 2,132 host species (Fig. 1c). Our pretraining corpus combines diverse microbial sequences with taxonomic annotations for taxonomy-informed pretraining.

**Figure 1.**
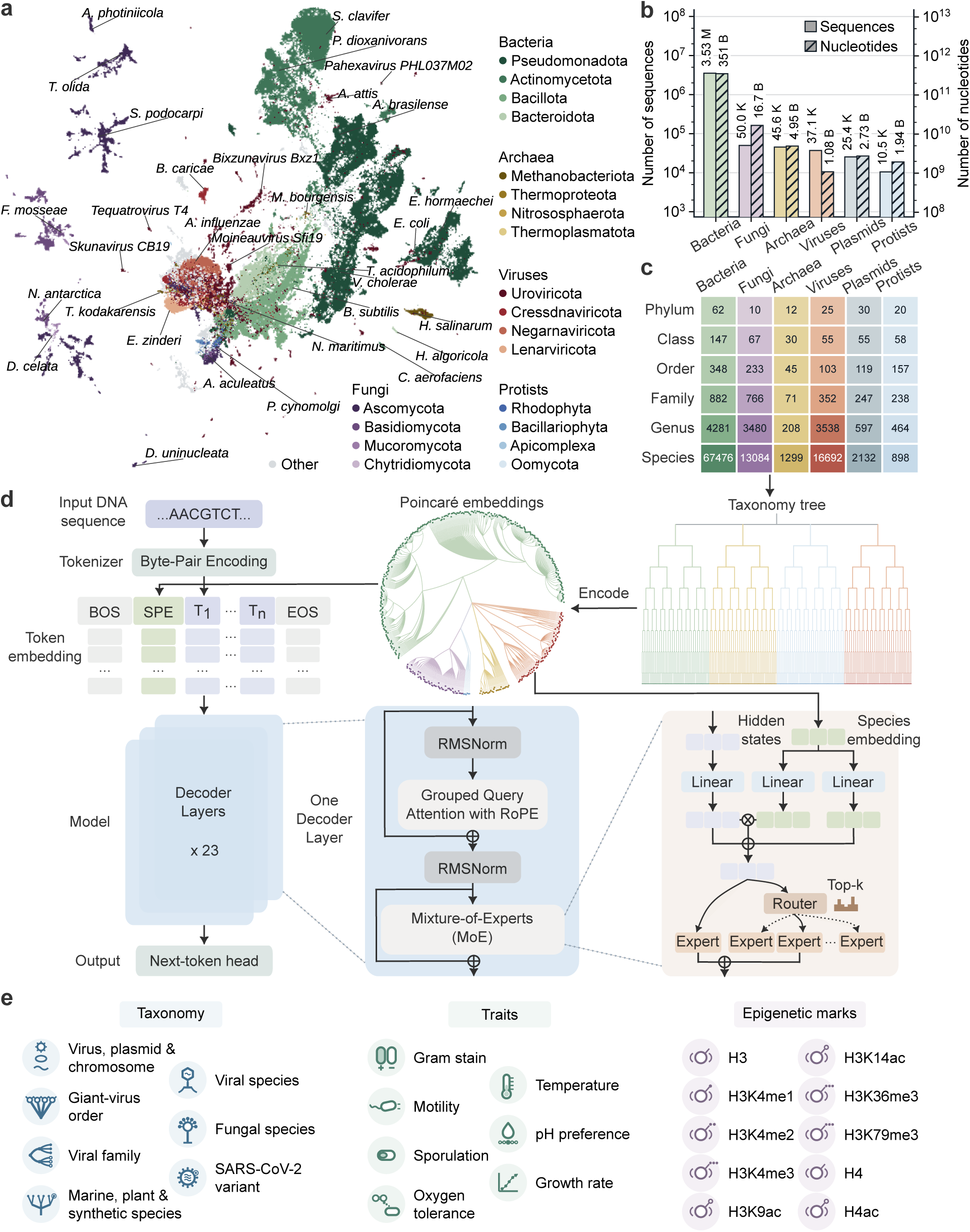
Overview of MicroGlot pretraining corpus, model architecture and evaluation tasks. **a**, Illustrative visualization of the pretraining corpus by uniform manifold approximation and projection (UMAP) ^34^ of *k*-mer frequency (*k* = 1 to 6). Each dot represents one of the 99,700 species. Colors distinguish the four phyla with the most species in each of the five labeled groups. Plasmid sequences carry host taxonomy, and are therefore not highlighted. Representative clusters are annotated with species names. **b**, Number of sequences and nucleotides for the six groups in the pretraining corpus. The y axes are on logarithmic scales. **c**, Number of distinct phyla, classes, orders, families, genera, and species within the six groups. Plasmids use host taxonomy and therefore may share labels with other groups. **d**, Overview of MicroGlot architecture. Poincaré embeddings ^30^ encode the corpus taxonomy, providing species embeddings for conditional sequence learning and guided expert routing. Decoder layers combine GQA ^31^, root mean square layer normalization (RMSNorm) ^35^, rotary position embeddings (RoPE) ^36^ and MoE modules for efficient language modeling. Beginning of sequence (BOS), species (SPE) and end of sequence (EOS) are special tokens. **e**, Evaluation tasks include microbial taxonomic classification, phenotypic trait prediction and epigenetic marks prediction.

### 2.2 MicroGlot incorporates taxonomic knowledge into a sparse mixture-of-experts architecture

Taxonomic classifications arrange organisms into taxa at successive ranks, providing relational information beyond species identity^26,37^. SpeciesLM ^28^ and SPACE^29^ introduce learned species embeddings, whereas Evo ^8^ and Evo2 ^18^ incorporate textual taxonomic lineages. Although these approaches supply biological context for conditional genomic sequence modeling, their embeddings are learned through prediction objectives without constraining embedding geometry to reflect the relational structures in taxonomy. To encode these structures, we adopted Poincaré embeddings^30^, whose hyperbolic geometry supports compact representations of hierarchical relationships. We learned these embeddings from the microbial taxonomy tree of our pretraining corpus, independent of any language modeling objective (Fig. 1d, Supplementary Fig. 2 and Methods). The encoded species embeddings serve as conditioning tokens and modulate expert routing through featurewise affine transformations^38^ (Methods), enabling sequence representation learning and expert selection to utilize the rich taxonomic knowledge present in established genome databases ^25,27,33^.

To combine this taxonomic context with scalable sequence modeling, we implemented MicroGlot as a decoder-only transformer pretrained by next-token prediction on DNA sequences tokenized with byte pair encoding (BPE). Sparse MoE layers ^23^ increase model capacity without requiring all experts to process every token. Each of the 23 decoder layers combines shared and selected experts, following residual MoE^39^. All tokens share one feedforward transformation, while the additional expert is selected according to sequence and taxonomic context (Methods). The backbone contains ~ 2.98 billion parameters but only sparsely activates 479 million per token, increasing model capacity to encompass the multilinguality of microbial genomics without introducing additional computational overhead.

Language specialization varies with depth in multilingual MoE models^40^, and allocating more experts to early and late layers has improved multilingual expansion^41^. With this motivation, we adopted a U-shaped allocation to provide greater capacity for taxon-dependent sequence patterns at these depths. We further limited model size by interleaving layers containing larger numbers of experts with layers containing only four experts (Methods). This follows the principle of alternating dense and sparse computation in Switch Transformer ^23^ and NLLB ^24^, while retaining conditional computation in every MicroGlot layer. Altogether, these designs make MicroGlot a taxonomy-informed DNA foundation model for multilingual representation learning in microbial genomics, combining hierarchical biological context with selective expert computation for microbial sequence analysis (Fig. 1e).

### 2.3 MicroGlot encodes microbial taxonomy and phenotypic traits in zero-shot embeddings from efficient pretraining

Genomic pretraining can yield representations that support biological prediction within a frozen backbone ^14,18^. We first asked whether MicroGlot’s zero-shot embeddings encoded information about both microbial taxonomic identity and phenotypic traits. Previous benchmarks have shown that probing performance varies across gLM layers, with the optimal depth depending on the specific model and task ^14,15,18^. Therefore, to ensure a fair comparison, we trained supervised probes on frozen embeddings at every layer and evaluated both layer-averaged and best-layer performance (Methods). We compared MicroGlot with multiple gLM baselines that contained microbial genomic data in their pretraining corpora, including ProkBERT-mini-long ^17^, DNABERT-2^12^, DNABERT-S^13^, three Nucleotide Transformer multispecies checkpoints^14^ and Evo2-7B-base ^18^. We evaluated MicroGlot on seven trait prediction tasks and six microbial taxonomic classification tasks (Table 1, Methods). For the taxonomic classification tasks, we used a separately pretrained variant, MicroGlot-plain, which is identical to MicroGlot in model and pretraining configurations but does not receive taxonomic knowledge, thereby isolating taxonomic information learned from the DNA sequence (Methods).

**Table 1.** Summary of benchmark datasets for layer-wise probing. The three replicon tasks use the same sequences with different labels.

| Task | Source | Samples | Classes |
| --- | --- | --- | --- |
| *Trait prediction* |  |  |  |
| Gram stain | BactoTraits<sup>42</sup> | 2,499 genomes | 2 |
| Motility | BactoTraits<sup>42</sup> | 2,357 genomes | 2 |
| Sporulation | BactoTraits<sup>42</sup> | 1,351 genomes | 2 |
| Oxygen tolerance | BactoTraits<sup>42</sup> | 2,103 genomes | 4 |
| Temperature | BactoTraits<sup>42</sup> | 2,899 genomes | 4 |
| pH preference | BactoTraits<sup>42</sup> | 890 genomes | 5 |
| Growth rate | Phydon<sup>43</sup> | 1,027 genomes | Regression |
| *Taxonomic classification* |  |  |  |
| Viral vs non-viral | geNomad<sup>44</sup> | 121,769 sequences | 2 |
| Plasmid vs non-plasmid | geNomad<sup>44</sup> | 121,769 sequences | 2 |
| Chromosome, plasmid & virus | geNomad<sup>44</sup> | 121,769 sequences | 3 |
| Giant-virus order | Global Ocean Eukaryotic Viral database<sup>45</sup> | 1,370 genomes | 4 |
| Viral family | Human gut virome study<sup>46</sup> | 473 sequences | 8 |
| Marine, plant & synthetic species | DNABERT-S benchmark<sup>13</sup> | 110,492 sequences | 1,208 |

To assess representation quality across model depth, we first evaluated layer-averaged probing performance. MicroGlot achieved the highest performance on all seven phenotypic trait prediction tasks, while MicroGlot-plain performed best on four of the six taxonomic classification tasks. Without taxonomic knowledge, MicroGlot-plain also outperformed all baselines on the seven trait prediction tasks (Supplementary Fig. 5a). Averaged across all 13 tasks, the two MicroGlot variants achieved a mean score of 0.816, compared with 0.798 for ProkBERT-mini-long, the top-performing baseline (Fig. 2a,b). We next evaluated each model at its best-performing layer, selected on the validation set (Methods). The two MicroGlot variants achieved either the highest or second-highest performance on every task within their respective evaluation sets. At these validation-selected layers, Evo2-7B-base achieved the highest mean score across tasks (0.841), followed by MicroGlot and MicroGlot-plain with a combined mean score of 0.835 (Fig. 2a,b). Together, these results show that MicroGlot learns representations that support both taxonomic classification and phenotypic trait prediction. Full layer-wise results are provided in Supplementary Figs. 5 and 6.

**Figure 2.**
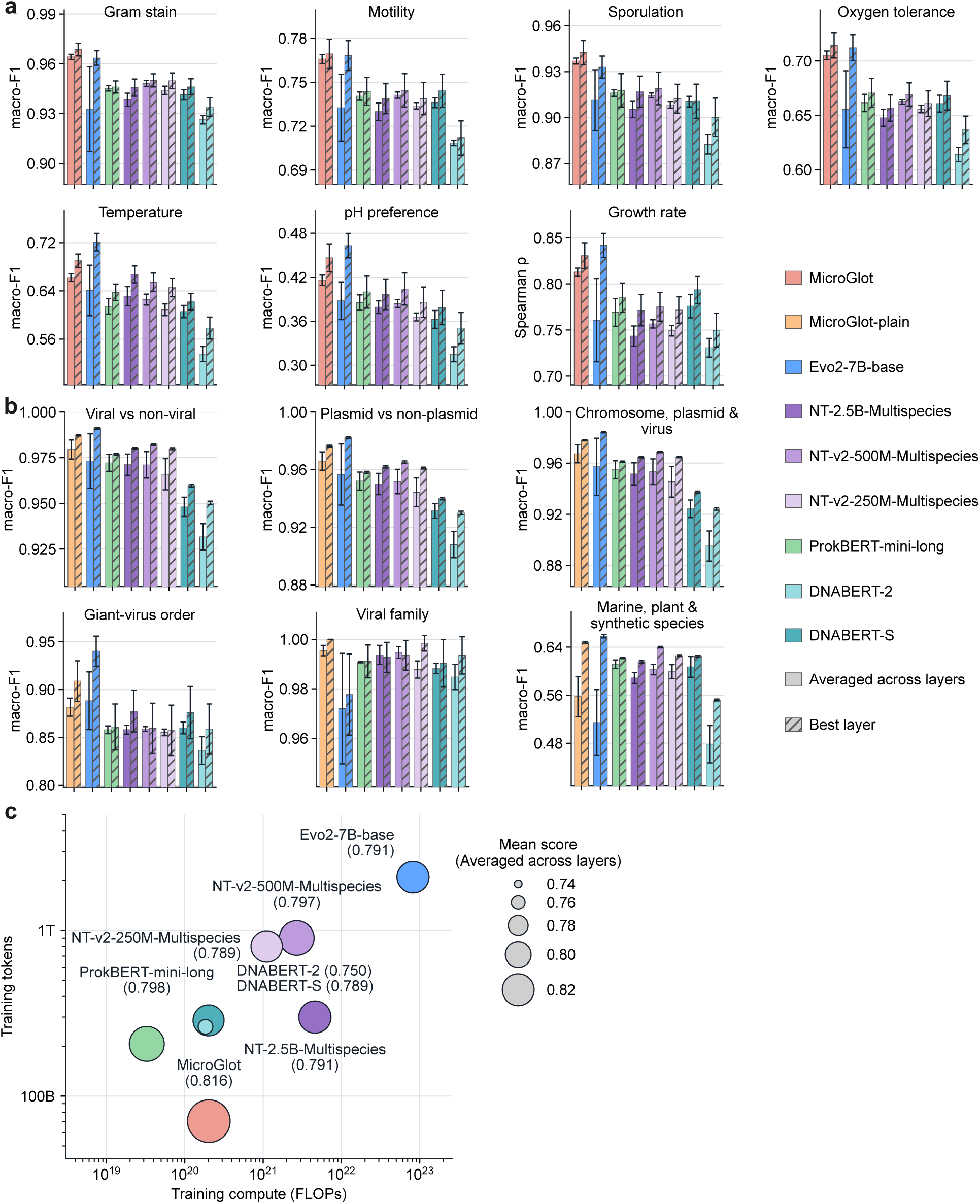
Benchmark results of layer-wise probing and pretraining compute. **a**, Benchmark results of MicroGlot and baseline gLMs on seven trait prediction tasks. **b**, Benchmark results of MicroGlot-plain with baseline gLMs on 6 microbial taxonomic classification tasks. In **a** and **b**, solid bars show scores averaged across layers and diagonally hatched bars show scores at the layer selected by validation performance. Error bars denote 95% confidence intervals calculated across layer means or across 10-fold cross-validation repeated with random seeds 0, 1 and 2, respectively. **c**, Pretraining token budgets and compute approximated by 6*ND* ^47^, with activated parameter count *N* and token count *D*. Baseline token budgets were obtained or derived from each model’s training description. Bubble size indicates the mean score across layers and tasks. The mean score for MicroGlot combines its seven trait scores with MicroGlot-plain’s six taxonomic classification scores.

MicroGlot learned high-quality representations using 70.6 billion pretraining tokens, fewer than any of the compared gLMs. With 479 million parameters activated per token, pretraining required approximately 2.03 10^20^ floating-point operations (FLOPs) (Fig. 2c). MicroGlot outperformed models with comparable total or activated parameter counts while requiring more than an order of magnitude fewer pretraining FLOPs. This computational efficiency is enabled by the sparse architecture, as top-1 expert routing restricts computation to only a small subset of parameters from the whole backbone. Modern language-model components such as GQA^31^, SwiGLU ^48^ feed-forward networks, and RMSNorm^35^ further contribute to computational efficiency. Together, these results demonstrate that MicroGlot learns informative representations of microbial taxonomy and phenotypic traits across model layers with comparatively low pretraining compute.

### 2.4 Taxonomic knowledge improves MicroGlot genomic representations

We assessed the contribution of taxonomic information to representation learning by comparing MicroGlot with MicroGlot-plain, an ablated variant that excluded the species token and species-guided routing. Both models used the same architecture and were pretrained for one epoch on the same corpus using an identical training scheme. Throughout pretraining, we observed that MicroGlot achieved lower validation perplexity at all 22 evaluation checkpoints than MicroGlot-plain (Fig. 3a, Methods).

**Figure 3.**
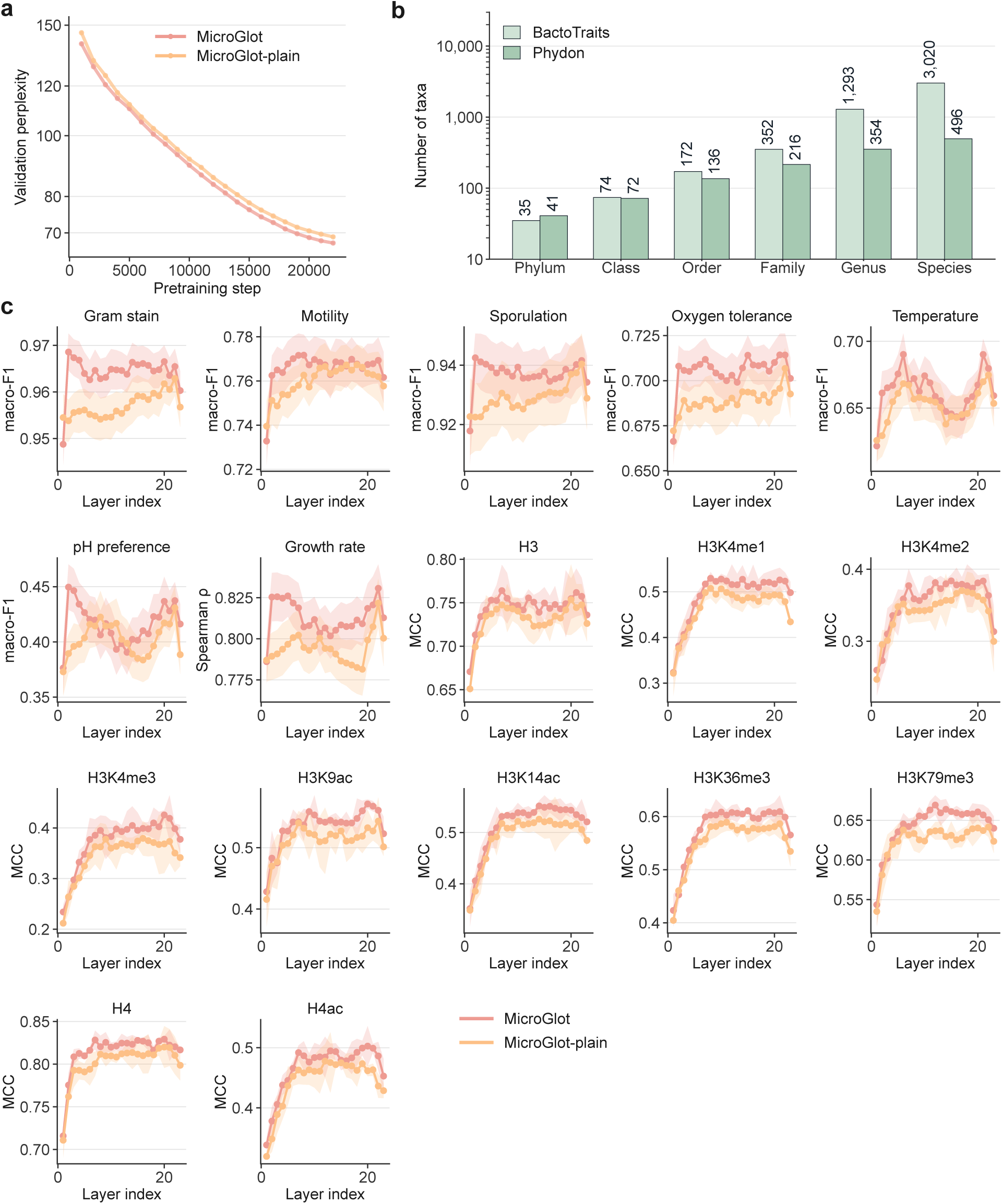
Ablation study of taxonomic knowledge in MicroGlot. **a**, Validation perplexity during one epoch of pretraining on the same corpus. MicroGlot-plain retains the backbone architecture but omits the species token and species-guided routing. Evaluation checkpoints were recorded every 1,000 steps. **b**, Numbers of distinct taxa from phylum to species in the 6 trait data from BactoTraits and the growth rate data from Phydon. Counts follow each dataset’s taxonomy and are shown on a logarithmic axis. **c**, Frozen probing performance across decoder layers 1 to 23 for seven traits and ten epigenetic marks. Lines show mean scores and shading denotes 95% confidence intervals. Trait tasks used 10-fold cross-validation with 3 random seeds. Epigenetic marks tasks used 3 random seeds with the fixed Genome Understanding Evaluation (GUE) ^12^ benchmark splits and the same *S. cerevisiae* species embedding for MicroGlot.

We next tested whether the advantage observed during pretraining translated to improved downstream biological prediction. We compared MicroGlot and MicroGlot-plain across the seven microbial trait prediction tasks and an additional ten epigenetic marks prediction tasks, probing representations from all 23 layers. The microbial trait datasets encompassed 3,020 species from 35 phyla in BactoTraits and 496 species from 41 phyla in Phydon, enabling evaluation across diverse microbial taxa (Fig. 3b). Species embeddings were retrieved from the precomputed reference set of 99,700 species (Methods). Matches were obtained for 2,848 of 3,020 (94.3%) BactoTraits species and 408 of 496 (82.3%) Phydon species, indicating broad coverage of the reference set. For species without a direct match, embeddings were inferred from DNA sequence using a separate species encoder (Methods). MicroGlot outperformed MicroGlot-plain in both best-layer and layer-averaged performance on all 17 tasks and in 370 of 391 task-layer comparisons across the 23 layers (Fig. 3c), demonstrating consistent benefit of taxonomy-informed pretraining across downstream tasks and model depth.

The epigenetic marks tasks provided a complementary within-species evaluation, in which all sequences were assigned the same *Saccharomyces cerevisiae* embedding. Despite this constant species-level input, MicroGlot achieved higher layer-averaged Matthews correlation coefficient (MCC) on all ten epigenetic marks, with the mean increasing from 0.531 to 0.552 (Fig. 3c). Because species identity did not vary across sequences, these gains cannot be attributed to differences in species identity among samples and instead indicate that taxonomy-informed pretraining improved the sequence representations themselves. Together, these results show that incorporating taxonomic information during pretraining improves the downstream utility of MicroGlot representations.

### 2.5 MicroGlot enables efficient genomic inference and fine-tuning

The practical value of a gLM also depends on the computational resources required for downstream use ^12,14^. We therefore examined whether MicroGlot’s sparse architecture could support efficient inference across increasing sequence lengths while maintaining competitive downstream performance at low fine-tuning cost. We benchmarked backbone inference across nine models spanning different architectural designs, including five encoder-based models, comprising ProkBERT-mini-long, DNABERT-2, and the NT and NT-v2 multispecies checkpoints, and four autoregressive models, comprising MicroGlot, Evo2-7B-1M, Evo2-20B-1M, and Evo2-40B-1M. The three Evo2 checkpoints use a StripedHyena convolutional architecture ^18^. For each model, we measured GPU memory use and inference latency on synthetic inputs at batch size one, with token lengths increasing in powers of two. At its trained context length of 8,192 tokens (~ 43.4 kb), MicroGlot required 6.27 GB of GPU memory and 80.6 ms per forward pass, substantially lower than baselines at the same context length (Fig. 4a,b). The efficiency advantage became more pronounced as sequence length increased. MicroGlot’s peak memory increased from 6.00 GB at 128 tokens to 6.27 GB at 8,192 tokens, corresponding to only a 4.5% increase over a 64-fold increase in context length. At twice the trained context length (~ 86.8 kb), MicroGlot required 6.54 GB and 167 ms, whereas the four encoder baselines that reached 80–98 kb required 35.9–55.0 GB and 0.94–4.17 s and exhausted available GPU memory at the next tested sequence length (Fig. 4a,b). At 131 kb (131,072 tokens for Evo2), Evo2-7B-1M required 90.69 GB and 15.3 s, whereas MicroGlot processed 694 kb (131,072 tokens) with 10.41 GB and 4.06 s (Fig. 4a,b). MicroGlot also reached a throughput of 538 kb/s at its trained context length, compared with 123–364 kb/s for the encoder baselines at their respective context lengths and at most 11.7 kb/s for the three Evo2 checkpoints at any tested length (Fig. 4b). Notably, MicroGlot further processed sequences of 2,097,152 tokens, equivalent to 11.1 Mb or 256 times its trained context length, using only 76.7 GB of GPU memory (Fig. 4a). By comparison, the longest sequences processed by the Evo2 models were 131 kb for Evo2-7B-1M, 32.8 kb for Evo2-20B-1M and 4.1 kb for Evo2-40B-1M. Together, these measurements show that MicroGlot maintains comparatively low memory use and inference latency as sequence length increases, supporting its potential for context extension.

**Figure 4.**
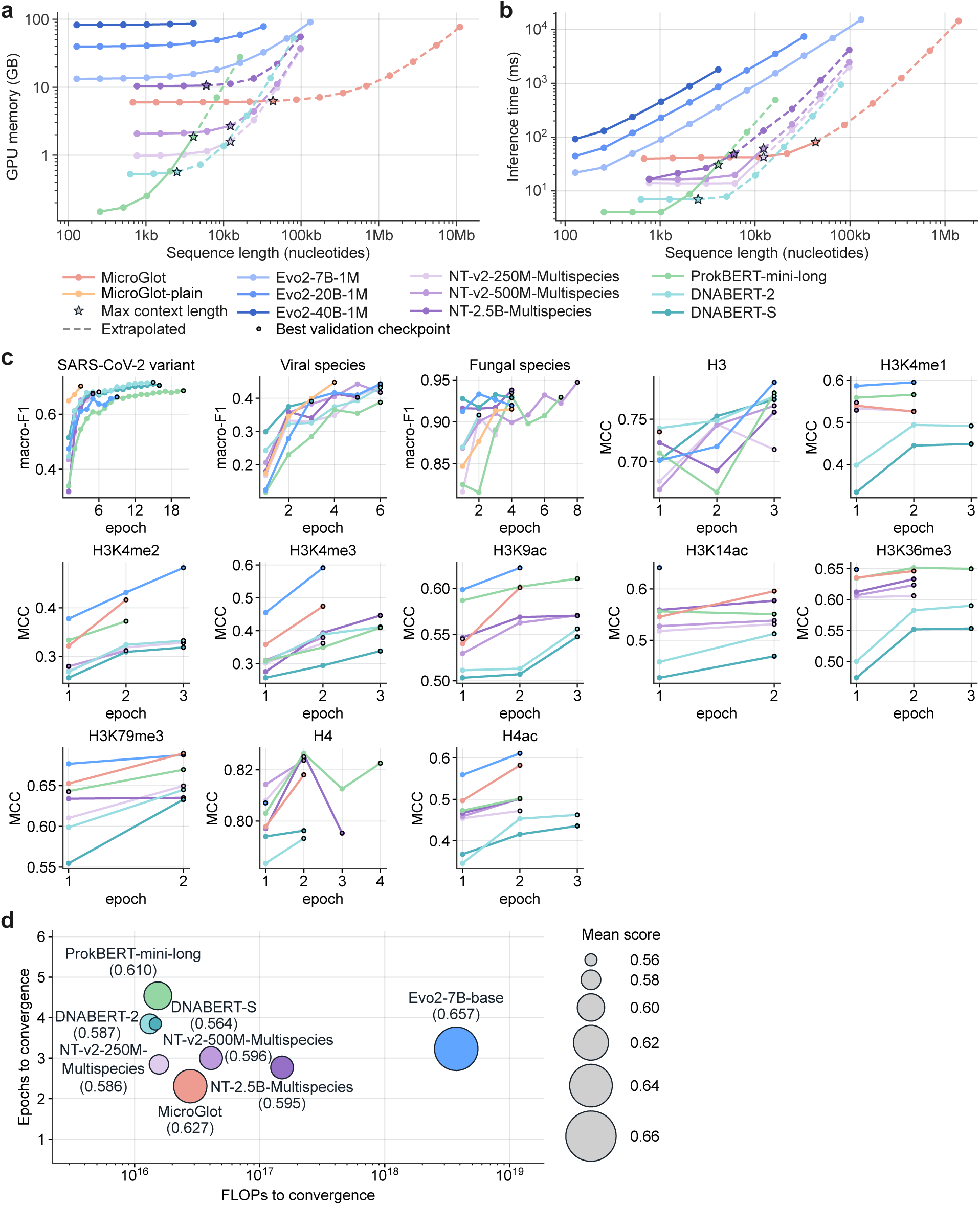
Comparison of computational efficiency in inference and fine-tuning. **a**, Peak allocated GPU memory across input lengths. **b**, Forward pass time across input lengths. Measurements in **a** and **b** timed backbone forward passes on synthetic inputs at batch size one. Each model was evaluated at context lengths doubling from 128 tokens until a single NVIDIA RTX PRO 6000 Blackwell Max-Q GPU ran out of memory. Stars mark configured context windows and dashed lines denote measured execution beyond context windows. Token counts were converted to kilobases using each model’s tokenizer compression ratio. **c**, Test scores on 13 microbial tasks from the GUE benchmark ^12^. Curves show individual fine-tuning runs and end at the checkpoint with the lowest validation loss, marked by a black-outlined circle. **d**, Mean epoch and compute at the selected checkpoint, with bubble size indicating the mean test score across the 13 tasks. The mean score for MicroGlot combines its epigenetic task scores with MicroGlot-plain’s taxonomic classification scores. Fine-tuning compute was accumulated over epochs through the selected checkpoint and averaged across tasks, approximated from each model’s parameter and processed token counts together with the quadratic cost of attention over the padded input lengths ^49^.

We next asked whether this computational efficiency could be retained during downstream adaptation without compromising predictive performance. We fine-tuned the models using Low-Rank Adaptation (LoRA) ^50^ on three microbial taxonomic classification tasks and ten epigenetic marks prediction tasks from the GUE benchmark^12^ (Methods, Supplementary Table 3 and Supplementary Table 4). Across the three taxonomic classification tasks, evaluated using macro F1, MicroGlot achieved the highest mean score among the eight evaluated models at 0.689, followed by Evo2-7B-base and DNABERT-2 at 0.686 and NT-v2-250M-Multispecies at 0.684 (Fig. 4c). MicroGlot also achieved the highest score on virus species classification, the most challenging of the three tasks by overall performance, with a macro F1 of 0.448 compared with 0.443 for Evo2-7B-base. Across all 13 tasks, MicroGlot achieved a mean score of 0.627, compared with 0.657 for Evo2-7B-base (Fig. 4c,d). This performance was reached after an average of 2.31 fine-tuning epochs, fewer than for any other evaluated model (Fig. 4c,d). MicroGlot required a mean of approximately 2.79 × 10^16^ FLOPs per task, compared with approximately 3.73 10^18^ FLOPs for Evo2-7B-base, corresponding to a ~ 134-fold difference in fine-tuning compute (Fig. 4d). MicroGlot also achieved a higher mean score with lower fine-tuning compute than the comparably sized baselines NT-2.5B-Multispecies (in total parameters) and NT-v2-500M-Multispecies (in activated parameters), whereas models requiring less compute achieved lower mean performance (Fig. 4d). Together, the inference and fine-tuning results show that MicroGlot combines competitive downstream performance with low computational requirements, supporting its practical use as a foundation model for microbial sequence analysis.

### 2.6 MicroGlot learns multilingual representations of microbial genomics through taxonomy-informed routing

Multilingual MoE models have been shown to share expert preferences across related languages^24^. We therefore investigated whether DNA sequences from related microbial taxa similarly exhibited shared expert-routing patterns in MicroGlot. To characterize expert usage across major taxonomic groups, we selected 1,000 species by balanced random sampling from the pretraining corpus (Methods), including 250 species from each of the three cellular microbial domains and 250 viruses. This balanced sampling yielded 3,948 sequence windows of 8 kb from 85 phyla, with 2–5 windows sampled per species, and enabled comparisons across taxonomically diverse groups (Methods).

Expert usage showed clear taxonomic structure across the 11 sparse layers. Different taxonomic groups exhibited distinct preferences for particular experts, although individual experts could be shared across multiple groups (Fig. 5a). To quantify these patterns, we defined a routing fingerprint for each species by calculating the fraction of DNA content tokens assigned to each expert within each window, concatenating these distributions across all 23 decoder layers, and averaging the resulting vectors across windows from the same species (Methods). Because tetranucleotide frequency (TNF) provides an established genomic signature for grouping microbial sequences ^51,52^, we used TNF profiles derived from the same windows as a biologically motivated reference. Across nine reference groups spanning bacteria, archaea, fungi, protists, and five viral realms, k-means clustering of routing fingerprints achieved a mean adjusted Rand index (ARI) of 0.620±0.056, compared with 0.115 ± 0.013 for TNF profiles over ten runs with different random seeds (Fig. 5b; Supplementary Note 5). Thus, routing fingerprints more closely recapitulated the reference taxonomic groups than TNF profiles.

**Figure 5.**
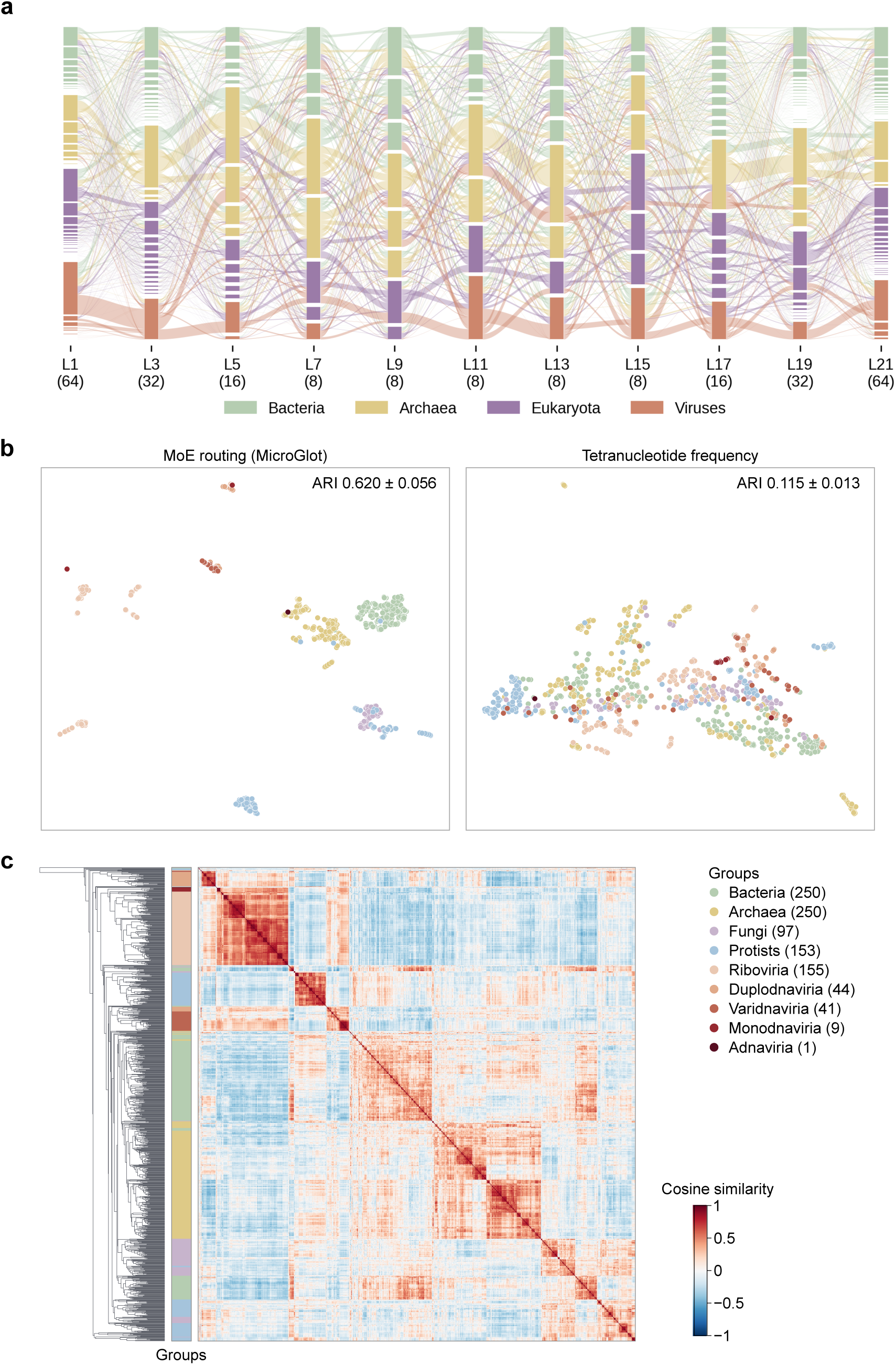
Taxonomic structures in MicroGlot routing fingerprints. **a**, Dominant expert assignments per window across the 11 layers containing more than 4 routed experts. Node heights and ribbon widths are proportional to window counts. Node color denotes the most frequent taxonomic group and ribbon color denotes the group of the contributing windows. **b**, UMAP of standardized species routing fingerprints and tetranucleotide frequency profiles, with one point per species. Colors distinguish bacteria, archaea, fungi, protists and five viral realms. ARI values are means ± standard deviations across 10 *k*-means runs on the standardized feature vectors, using 9 clusters. **c**, Cosine similarity between routing fingerprints of the 1,000 species. Numbers in the shared legend of **b** and **c** show species counts.

This taxonomic organization was also apparent in pairwise cosine similarities between species-level routing fingerprints. Mean similarity was 0.295 for species within the same reference group and similar routing patterns (Fig. 5c). These results indicate that sequences from the same broad taxonomic group tend to recruit more similar combinations of experts, while expert usage remains shared across taxa rather than being restricted to individual groups. Together, these analyses reveal a taxonomically structured organization of expert routing in MicroGlot. Combined with the representation gains observed in the ablation study (Fig. 3), these results suggest that taxonomic information shapes not only the learned genomic representations but also how taxonomic signals are routed through the model architecture.

## 3 Discussion

We pretrained MicroGlot, a taxonomy-informed gLM designed for multilingual representation learning from microbial DNA. MicroGlot encodes taxonomic relationships using Poincaré embeddings ^30^ and integrates this knowledge into a sparse MoE architecture to condition both sequence representation learning and expert routing. Across the evaluated gLMs, MicroGlot achieved the highest layer-wise probing performance while using the smallest pretraining token budget (Fig. 2). Ablation experiments further showed that incorporating taxonomic information improved learned representations across model depth (Fig. 3), while downstream fine-tuning demonstrated competitive predictive performance at substantially lower computational cost (Fig. 4). Analysis of expert routing further revealed taxonomically structured routing patterns, with species-level routing fingerprints recapitulating taxonomic groups more closely than tetranucleotide frequency profiles (Fig. 5). Together, these findings show that explicitly incorporating taxonomic knowledge can improve microbial genomic representation learning while retaining the computational advantages of sparse modeling. Several directions could further extend the scope of MicroGlot. Our inference benchmarks showed comparatively modest growth in GPU memory as sequence length increased and lower latency than the evaluated baselines at comparable long input lengths, suggesting that the architecture could support further extension of its trained context (Fig. 4a,b). Evo^8^, Evo2 ^18^, and Genos ^53^ extended their context lengths through staged pretraining on progressively longer sequences, with Evo2 and Genos reaching approximately one million nucleotides. Applying a similar training strategy to MicroGlot could enable modeling of longer-range genomic dependencies, including distal regulatory interactions. Taxonomic coverage could likewise be expanded. The current reference set contains precomputed embeddings for 99,700 species spanning major microbial cellular domains and viral realms (Fig. 1). By comparison, SpeciesLM^28^ represents 806 fungal species, whereas SPACE^29^ incorporates species information from *Homo sapiens* and *Mus musculus*. Despite its broader microbial coverage, MicroGlot still represents only a fraction of the estimated global diversity of microorganisms ^54^. Continued expansion of RefSeq^25^ and ongoing revisions to ICTV taxonomy^27^ provide opportunities to incorporate newly characterized taxa and updated taxonomic relationships. Expanding both the pretraining corpus and the species embedding reference set could therefore extend taxonomy-informed modeling to a broader range of microbial diversity.

## 4 Methods

### 4.1 Pretraining corpus

We constructed the pretraining corpus from RefSeq^25^ (release 232), the ICTV Virus Metadata Resource ^27^ (VMR; release MSL40 v1) and PLSDB^33^ (version 2024_05_31_v2). The collection comprised bacterial, archaeal, fungal, protist and viral sequences, together with plasmids. We assigned NCBI Taxonomy^26^ lineages to RefSeq sequences and PLSDB hosts and obtained ICTV lineages from the VMR. These annotations span eight ranks, including cellular domain or viral realm, kingdom, phylum, class, order, family, genus and species. Plasmid sequences retained the taxonomy of their annotated hosts.

The combined collection comprised 46,733,579 sequences totaling 2.00 trillion nucleotides. We converted raw sequences to uppercase and trimmed leading and trailing ambiguous bases. Here, ambiguous bases were defined as any character other than A, C, G or T, including N and the other IUPAC ambiguity codes ^55^. We excluded the 140,466 sequences (0.30%) in which these characters constituted more than 5% of the trimmed sequence and replaced the remaining ambiguous bases with N. This left 46,593,113 sequences totaling 1.99 trillion nucleotides. We represented the lineage of each sequence as a root-to-leaf path through the eight ranks, treated missing ranks as gaps in this path and imputed these gaps using a series of taxonomy-based rules (Supplementary Note 1). Most gaps occur where the source taxonomies assign no taxon at a rank, such as the kingdom of protists or the order of most tailed phages, and are therefore concentrated in particular clades. We kept sequences of at least 500 bp with annotations at all eight taxonomic ranks, removing 8,409,770 shorter sequences and a further 19,777 whose annotations remained incomplete after imputation. This left 38,163,566 sequences, including 459,502 (1.20%) that would have been excluded without imputation. Across the 99,700 species of the final corpus, imputation filled 19,027 of the 797,600 rank entries (2.39%). We then randomly sampled up to 100 sequences per species, retaining all sequences for species with no more than this number. This yielded 3,725,357 sequences, among which 30,292 were further removed due to duplicated accessions in more than one database. The final pretraining corpus contained 3,695,065 sequences spanning 378.3 billion nucleotides across 99,700 species (Fig. 1b,c).

### 4.2 Tokenization

We chose byte pair encoding (BPE)^56^, which merges frequent adjacent segments into variable-length tokens. BPE compresses sequences several-fold relative to single-nucleotide or overlapping *k*-mer tokenization, allowing a fixed token window to span longer genomic context. We trained the tokenizer using one sequence sampled uniformly at random from each species in the pretraining corpus. For sequences longer than 100 kb, we sampled a single contiguous 100 kb window at a random position. This yielded 4.23 billion nucleotides for tokenizer training. We used the BPE implementation in the Hugging Face Tokenizers library, with a minimum merge frequency of zero and no maximum token length. We split sequences at N to prevent merges across ambiguous positions.

To determine the optimal vocabulary size, we compared vocabularies containing 4,096, 8,192, 16,384 and 32,768 tokens using 30,000 sequences sampled uniformly at random, with equal numbers drawn from each of the bacteria, archaea, fungi, protists, viruses and plasmids groups. We measured sequence compression as nucleotides per token and vocabulary utilization as the fraction of tokens occurring at least 100 times within each group. We selected a vocabulary of 8,192 tokens, which had an average of 5.32 nucleotides per token and at least 99.7% utilization in every group. Vocabularies of 16,384 and 32,768 tokens provided modest gains in compression ratio but reduced minimum utilization to 91.4% and 67.0%, respectively (Supplementary Fig. 1). Vocabulary tokens ranged from 1 to 16 nucleotides in length (median 7), and across the full corpus the trained tokenizer encoded an average of 5.42 nucleotides per token. Tokenized sequences consisted of a beginning-of-sequence (BOS) token, DNA content tokens, an end-of-sequence (EOS) token and padding (PAD) tokens where required, with N representing ambiguous bases.

### 4.3 Species embeddings

We constructed a taxonomic hierarchy by linking each taxon to its parent at the rank above. Above the domain or realm rank, we linked the three domains (i.e., Bacteria, Archaea and Eukaryota) to a node for cellular microorganisms and the seven realms (i.e., Adnaviria, Duplodnaviria, Monodnaviria, Riboviria, Ribozyviria, Singelaviria and Varidnaviria) to a node for viruses, and joined both to a single root node, following NCBI Taxonomy. We then computed the transitive closure, comprising all ancestor-descendant pairs along each lineage. We applied the Poincaré embedding algorithm^30^ to this transitive closure to encode an embedding for each node, using the reference implementation at https://github.com/facebookresearch/poincare-embeddings. Embeddings were optimized by sparse Riemannian stochastic gradient descent^57^ for 1,500 epochs, with a batch size of 1,024, 50 negative samples and a base learning rate of 0.02.

We evaluated embedding dimensions of 2, 4, 8, 16, 32, 64, 128, 256, 512 and 1,024 using two criteria. First, we evaluated retrieval of taxonomic relationships ranked by Poincaré distance using mean average precision (MAP) ^30^. Second, we trained multinomial logistic regression probes to predict order, family and genus from the embeddings and evaluated their accuracy using 5-fold cross-validation. We selected 32 dimensions, which yielded a MAP of 0.869, and probe accuracies of 0.934, 0.778 and 0.543 across 485 orders, 1,160 families and 2,719 genera, respectively. Increasing the dimensionality beyond 32 provided no substantial improvement (Supplementary Fig. 2). Supplementary Note 2 describes the Poincaré distance, the training objective and the calculation of MAP.

### 4.4 Model architecture

Here we describe MicroGlot’s architecture, with more detailed formulations deferred to Supplementary Note 3. Key architectural specifications are summarized in Supplementary Table 1. MicroGlot is a decoder-only MoE-based transformer with 23 layers. Each layer comprises an attention sublayer and an MoE sublayer, each preceded by RMSNorm^35^ and followed by a residual connection. For layer *l* = 1*, . . . ,* 23 and the *T* positions *t* = 0*, . . . , T* − 1 of an input window,

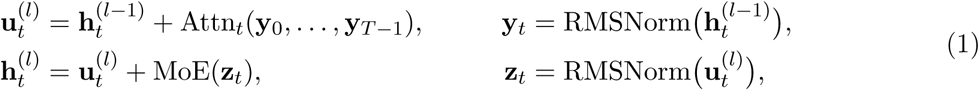

where 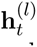 is the hidden state at position *t* after layer *l*, 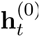 is the input embedding and Attn*_t_* is the causal self-attention output at position *t*. Attention uses GQA^31^ with 16 query heads and 8 key and value heads, and encodes token positions using RoPE^36^ with a base of 500,000. Attention was implemented using PyTorch scaled dot-product attention with the FlashAttention-2 backend^32^.

Each MoE sublayer combines a shared expert FFN^(s)^ with one of *E_l_* routed experts FFN_*i*_^(r)^ (*i* = 1*, . . . , E_l_*), selected by top-1 routing, following the residual MoE architecture^39^. All experts are SwiGLU ^48^ feed-forward networks. A router **W**_r_ maps the MoE input **z** (indices *l* and *t* omitted) to logits **W**_r_**z** over the routed experts, which are modulated by the species embedding and converted by a softmax to routing probabilities **p** = (*p*_1_*, . . . , p_E_* )^⊤^ as given in equation (3). The output of the selected expert *i*^∗^ = arg max*_i_ p_i_* is combined with the shared-expert output by per-token mixing weights,

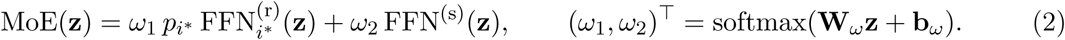

The number of routed experts *E_l_* was allocated across model depth in a U-shaped pattern (Supplementary Fig. 3). The 11 sparse layers contained 64, 32, 16, 8, 8, 8, 8, 8, 16, 32 and 64 experts, respectively, whereas all remaining layers contained four. This allocation was motivated by evidence of greater language specificity near the input and output layers of multilingual dense ^58,59,60^ and MoE ^40,41,61^ models, and by large MoE language models that interleave sparse layers with relatively more dense layers ^23,62,63^. This design increases model capacity for multilingual representation learning while keeping the computational cost manageable.

Taxonomic information is incorporated through an input token and expert routing. A learned linear projection **W**_spe_ maps the species embedding, normalized to unit Euclidean norm and denoted **e***_s_ ∈* R^32^, to a species (SPE) token 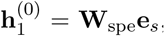, inserted after the BOS token. The SPE token participates in causal attention, so that all subsequent tokens can attend to it, but is excluded from representation pooling. In each MoE sublayer, feature-wise linear modulation (FiLM)^38^ scales and shifts the routing logits before the softmax,

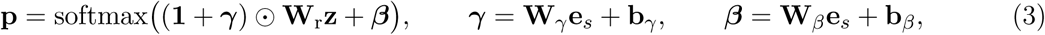

where **1** is the all-ones vector, denotes elementwise multiplication and **W***_γ_,* **b***_γ_,* **W***_β_,* **b***_β_* are learned for each layer.

### 4.5 Pretraining

We created a species-stratified validation set by randomly holding out one sequence from each species with more than ten sequences, resulting in 54,779 sequences (~ 1.5% of the pretraining corpus). Sequences longer than 8,190 tokens were divided into windows of 8,190 tokens, with an overlap of 200 tokens between consecutive windows, and the final window of each sequence contained its remaining tokens and was therefore usually shorter. Shorter sequences formed single windows. Each window was flanked by BOS and EOS tokens.

MicroGlot was pretrained by next-token prediction loss, combined with an auxiliary load-balancing loss ^23^ applied at every MoE layer to encourage a balanced load across experts. Without this loss, learned gating tends to route most tokens to a small number of experts, leaving the remaining experts insufficiently trained^62^. The total loss was

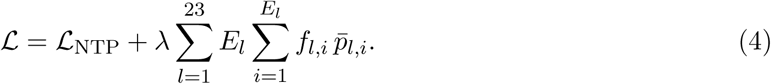

Here L_NTP_ is the mean next-token prediction cross-entropy over sequence tokens, excluding N, BOS, SPE and the padding token. For layer *l*, *f_l,i_* is the fraction of positions in the batch routed to expert *i* under top-1 selection, and *p̂_l,i_* is the routing probability *p_i_* of expert *i* (equation (3)) averaged over the same positions. The coefficient *λ* was set to 0.001 (Supplementary Table 1). For the taxonomic ablation, MicroGlot-plain omitted the SPE token and FiLM (***γ*** = ***β*** = **0** in equation (3)), with all other architectural and training settings unchanged.

We pretrained the model using AdamW with *β*_1_ = 0.9, *β*_2_ = 0.95, *ɛ* = 10^−8^, weight decay 0.1 and gradient clipping at a norm of 1.0. Following a warmup over the first 1% of training steps, the learning rate decreased from 3 x 10^−4^ to 3 x 10^−5^ according to a cosine schedule. The global batch size was ~ 4 million tokens per optimizer step. Training used the Transformers (v.4.51.3) model and training interfaces with bfloat16 precision, gradient checkpointing and DeepSpeed (v.0.18.4) ZeRO stage 1 for distributed training. Routing probabilities and auxiliary losses were computed in float32 (Supplementary Table 2).

We trained a separate species encoder to predict species embeddings from DNA sequences when direct lookup in the reference set of 99,700 species was unavailable. The species encoder used the same architecture and pretraining scheme as MicroGlot (Supplementary Table 1 and Supplementary Table 2). The final EOS representation was passed through a projection head comprising a 1,024-unit hidden layer, GELU ^64^ activation and a 32-dimensional output. The species encoder was trained with mean squared error loss to predict the normalized reference species embeddings, with the load balancing term also included. Training settings were identical to MicroGlot pretraining. The species encoder achieved a mean cosine similarity of 0.9744 between predicted and reference embeddings on the held-out validation set.

MicroGlot, MicroGlot-plain and the species encoder were each trained for a single epoch on the same corpus with 8 NVIDIA RTX PRO 6000 Blackwell GPUs for ~ 5 days. Each training run comprised 22,087 optimizer steps and processed 70.56 billion tokens excluding padding, or 91.19 billion tokens after padding each window to the 8,192-token context length.

### 4.6 Benchmark datasets

We evaluated the models on 13 probing tasks and 13 fine-tuning tasks (Table 1 and Supplementary Table 4). The six taxonomic probing tasks covered replicon type (geNomad ^44^), viral family ^46^ and giant virus order ^45^, both distributed with Virgo^65^, and species classification in marine, plant and synthetic species (DNABERT-S benchmark ^13^ from CAMI II^66^ and reference genomes). The seven trait probing tasks used species-level BactoTraits annotations ^42^ matched to RefSeq assemblies ^25^ and Phydon minimum doubling times ^43^, modeled on a base-two logarithmic scale. Each probing dataset was deduplicated at 90% nucleotide identity. The fine-tuning datasets were taken from GUE^12^ and used with their supplied splits, without additional deduplication. Details of benchmark dataset processing are provided in Supplementary Note 4, and sequence length distributions of the probing datasets are shown in Supplementary Fig. 4.

### 4.7 Probing

We evaluated frozen representations from each model layer using supervised probes, following the layer-wise evaluation of NT ^14^. Baselines included ProkBERT-mini-long ^17^, DNABERT-2^12^, DNABERT-S^13^, three NT multispecies checkpoints^14^ (NT-v2-250M-Multispecies, NT-v2-500M-Multispecies and NT-2.5B-Multispecies) and Evo2-7B-base^18^. From each layer, we extracted token embeddings and obtained a sequence embedding by mean pooling. Padding tokens were excluded from pooling for all models, and SPE tokens were excluded for MicroGlot. Sequences longer than a model’s context length were split into consecutive non-overlapping windows, embeddings were extracted from each window separately, and the window embeddings were averaged to obtain one embedding per sequence, following HyenaDNA ^11^. Giant virus order classification and the seven trait tasks used genome assembly-level labels. For these tasks, embeddings were extracted from each contig separately, and the contig embeddings were averaged to obtain one embedding per assembly. The remaining five tasks classified individual sequences (Supplementary Note 4).

We used 10-fold cross-validation with training, validation, and test sets split in an 8:1:1 ratio. Each experiment was repeated 3 times with seeds 0, 1 and 2. Epigenetic marks probing in the ablation study instead retained the predefined GUE benchmark splits. For each training split, we standardized the embeddings and trained a multilayer perceptron with a 1,024-unit hidden layer, ReLU activation and a classification or regression output. Probes were optimized using AdamW with default moment parameters, a constant learning rate of 10^−4^, weight decay 0.01, gradient clipping at a norm of 1.0 and a batch size of 64. Probes were trained for at most 100 epochs, with early stopping after 5 epochs without improvement in validation loss. For each model and task, we selected the layer with the highest mean validation score and reported its test performance.

### 4.8 Fine-tuning

We fine-tuned all models with LoRA^50^, training the LoRA parameters and a classification head while keeping the pretrained backbone frozen. Using the parameter-efficient fine-tuning (PEFT) library (v.0.13.2), we applied LoRA to all linear layers of each backbone with rank *r* = 16, *α* = 32 and dropout 0.05, keeping all other LoRA settings at their defaults. The ratio *α/r* = 2 and the dropout follow the NT configuration used in DNABERT-2^12^. Training used the Trainer of Hugging Face Transformers (v.4.51.3) with AdamW, a batch size of 32, a weight decay of 0.01 and gradient clipping at a norm of 1.0. The learning rate increased linearly over 500 warmup steps to 10^−4^ and then followed a cosine decay to 10^−5^. Models were trained for up to 100 epochs with early stopping after three epochs without improvement in validation loss, and the checkpoint with the lowest validation loss was evaluated on the test set (Supplementary Table 3).

Sequence representations were obtained by mean pooling the final-layer token embeddings, excluding padding tokens for all models and SPE tokens for MicroGlot. These representations were passed to a classification head consisting of a fully connected hidden layer with 1,024 units and ReLU activation, followed by a linear output layer. Sequences exceeding each model’s context length were chunked into non-overlapping windows. During training, each window was assigned the label of its source sequence and treated as a separate training example. At test time, class probabilities were averaged across windows to obtain a single prediction for each original sequence.

### 4.9 Sequence composition and routing fingerprints

For the pretraining corpus visualization (Fig. 1a), we represented each of the 99,700 species by *k*-mer composition, computed from its training sequences as the frequencies of *k*-mers for *k* = 1 to 6 concatenated into a feature vector. Ambiguous nucleotides (N) were skipped. These vectors were standardized, reduced by principal component analysis (PCA) (100 principal components) and visualized using UMAP.

For routing analysis, we selected 1,000 species via taxonomically stratified sampling, drawing 250 species each from Bacteria, Archaea, Eukaryota and Viruses. For every species, we randomly sampled up to five distinct sequences and took the central 8,000-nucleotide window of each, yielding 3,948 windows. For each window and decoder layer, we computed the fraction of DNA content tokens that were assigned to the selected expert. Concatenating these fractions over the 23 decoder layers yielded a 312-dimensional routing fingerprint per window. In the routing path visualization (Fig. 5a), each window was assigned to the expert selected most frequently among its DNA content tokens. Further details are provided in Supplementary Note 5.

## Data availability

Pretraining sequences are available from NCBI RefSeq ^25^ release 232, whose sequence accessions are listed in the release catalog at https://ftp.ncbi.nlm.nih.gov/refseq/release/release-catalog/archive/RefSeq-release232.catalog.gz, the ICTV Virus Metadata Resource ^27^ (VMR) release MSL40 v1, dated 7 March 2025, at https://ictv.global/sites/default/files/VMR/VMR_MSL40.v1.20250307.xlsx, and PLSDB^33^ version 2024_05_31_v2, whose archive version 2 was published on 7 January 2025, at https://doi.org/10.6084/m9.figshare.27252609.v2. Sequences specified by ICTV accessions are available from NCBI Nucleotide at https://www.ncbi.nlm.nih.gov/nuccore/. Pretraining and benchmark taxonomic annotation used the NCBI Taxonomy ^26^ snapshot of 1 September 2025 at https://ftp.ncbi.nlm.nih.gov/pub/taxonomy/taxdump_archive/new_taxdump_2025-09-01.zip. Sources for the benchmark datasets are given in Supplementary Note 4.

## Code availability

MicroGlot is available on Hugging Face at https://huggingface.co/athanzli/MicroGlot. Source code is available on GitHub at https://github.com/athanzli/MicroGlot.

## Acknowledgements

None.

## Author contributions

R.L., Y.D. and A.Z.L. conceived the research idea. R.L. and Y.D. supervised the study. A.Z.L. conducted the literature review. A.Z.L. and S.W. curated the datasets. A.Z.L. and S.C. designed the methodology and implemented the model. A.Z.L. trained the model and conducted the experiments. A.Z.L. analyzed the results, generated the visualizations and drafted the manuscript.

## Supplementary Material

### Supplementary Notes

**Supplementary Note 1. Imputation of missing taxonomic ranks**

We assigned each sequence a lineage over eight ranks (domain or realm, kingdom, phylum, class, order, family, genus and species) using NCBI Taxonomy for RefSeq sequences and the hosts of PLSDB plasmids, and the VMR for ICTV sequences. Some of these ranks had no label in the source annotations, mostly because the source taxonomies do not assign a taxon at every rank. For example, NCBI Taxonomy assigns no kingdom to protist lineages, most species of the class *Caudoviricetes* have no order, and the realm *Ribozyviria* has no taxon between realm and family. The gaps are therefore concentrated in particular clades, and excluding incomplete lineages would have removed entire groups, including all 898 protist species, all 21 *Ribozyviria* species and more than a third of the 16,692 virus species. We therefore filled these gaps with three rules, applied in decreasing order of evidence to the lineages of all 46,593,113 sequences that remained after removing sequences with more than 5% ambiguous bases, before length filtering and per-species sampling. Imputation also placed every species at the same depth of the taxonomic hierarchy to enable taxonomic encoding using the Poincaré embedding algorithm (Supplementary Note 2). Each rule acted only on entries left missing by the preceding rules. For the 99,700 species of the pretraining corpus, we counted a rank entry as imputed when none of the species’ pretraining sequences carried a label at that rank in the source annotations; 19,027 of the 797,600 entries (2.39%) were imputed, and the counts given below for each rule refer to these entries.

Rule one used the NCBI Taxonomy tree. RefSeq sequences carry an NCBI taxon identifier from the RefSeq release catalog, and PLSDB plasmids carry the identifier of their host taxon. Starting from this identifier, we followed the parent taxa up the NCBI Taxonomy tree to the root and filled each missing rank with the name of the ancestor at that rank. For example, a PLSDB plasmid of *Acidovorax* sp. JS42 (taxon identifier 232721) had no kingdom label, and the kingdom among its ancestors, *Pseudomonadati*, was assigned. In practice, this rule completed only PLSDB lineages, mainly by adding the kingdom rank that PLSDB host taxonomy lacks, because RefSeq lineages were derived from the same tree. Sequences from the ICTV Virus Metadata Resource carry no NCBI taxon identifier, so this rule did not apply to them. Rule one resolved 232 entries.

Rule two used the labels of related sequences. Sequences were grouped by a lower-rank taxon, and a missing label at a higher rank was filled when all sequences in the group that carried that label agreed on it. If they disagreed, the entry remained missing. Groups were formed at each rank from species up to kingdom. For example, NCBI Taxonomy places the genus *Rhodobacter* directly under the order *Rhodobacterales*, without a family, so its RefSeq sequences lacked a family label, whereas the seven *Rhodobacter* plasmids from PLSDB all listed *Paracoccaceae* in their host annotation. The missing family labels were therefore set to *Paracoccaceae*. Rule two resolved 17 entries, all of which were this family label for 17 *Rhodobacter* species.

Rule three filled remaining missing entries with a synthetic placeholder label. A placeholder stands for an unnamed taxon at the missing rank within its parent taxon and is formed by joining the initial of the missing rank (k, p, c, o, f, g or s) to the label of the parent with underscores. For instance, bacteriophage *λ* (*Lambdavirus lambda*) belongs to the class *Caudoviricetes* and the genus *Lambdavirus* but has no assigned order or family. It therefore received the order o Caudoviricetes, denoting an unnamed order within *Caudoviricetes*, and the family f o Caudoviricetes, denoting an unnamed family within that order. A placeholder thus adds no information beyond the known ancestor and never conflicts with an existing label, while completing the lineage of the sequence. Sequences lacking a label under the same named ancestor receive the same placeholder and are treated as members of one unnamed taxon. Rule three resolved the remaining 18,778 entries. Most of these (14,635; 77.9%) were in viral lineages, mainly the orders and families of tailed phages (class *Caudoviricetes*; 7,815 entries) and the ranks from kingdom to genus of 948 RNA virus species assigned only to the realm *Riboviria* (5,688 entries). Among cellular organisms, the largest group was the kingdom of all 898 protist species.

**Supplementary Note 2. Poincaré embeddings and evaluation**

Poincaré embeddings ^30^ represent each node of the taxonomic hierarchy by a point **x** inside the unit ball, that is, a vector with Euclidean norm ‖ x ‖ *<* 1. In this Poincaré ball model of hyperbolic space ^30,67^, the Riemannian metric scales Euclidean lengths at **x** by the factor 2*/*(1 - ‖**x‖** ^2^), which gives the Poincaré distance between nodes *u* and *v* with embeddings **u** and **v**

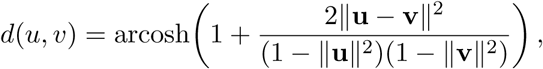

where arcosh is the inverse hyperbolic cosine. Because distances grow rapidly toward the boundary, general taxa can lie near the origin and their descendants near the boundary, far apart from one another ^30^.

The embeddings were learned from the set D of ordered pairs (*u, v*) in which *v* is an ancestor of *u*, which form the transitive closure. Following Nickel and Kiela ^30^, we minimized

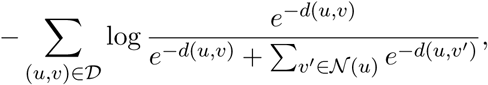

where N (*u*) contains negative samples drawn for each pair from the nodes that are neither *u* nor its ancestors. The loss thus favors shorter distances to ancestors than to sampled nodes.

We evaluated how well the distances reconstruct the hierarchy, following the evaluation approach adopted by Nickel and Kiela ^30^. Each node *u* except the root, which has no ancestors, served as a query, and the remaining nodes, from all taxonomic ranks, were ordered by increasing Poincaré distance from *u*. The *m_u_* ancestors of *u* (nine for each species, including its seven higher-rank taxa, the cellular or virus node and the root node) were relevant, and all other nodes, including the descendants of *u*, were non-relevant. Average precision (AP) was defined as in information retrieval ^68^,

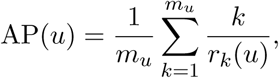

where *r_k_*(*u*) is the position of the *k*th closest ancestor of *u* in this ordering, so that *k/r_k_*(*u*) is the precision within the top *r_k_*(*u*) nodes. AP equals 1 when all ancestors of *u* are closer to it than any other node. Mean average precision (MAP) is the mean of AP over all query nodes.

**Supplementary Note 3. Model architecture details**

MicroGlot is a decoder-only transformer with *L* = 23 layers of hidden dimension *d* = 1,024, *H* = 16 query heads and *G* = 8 key and value heads of dimension *d*_h_ = *d/H* = 64, experts of intermediate dimension *d*_ff_ = 2,816, a vocabulary ν of │ν│ = 8,192 tokens and species embeddings of dimension *d*_s_ = 32 (Supplementary Table 1).

### Input sequence

An input window (*x*_1_*, . . . , x_n_*) of tokens *x_t_* ∈ ν begins with the BOS token *x*_1_. The species embedding of the sequence (Supplementary Note 2), normalized to unit Euclidean norm and denoted 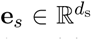, is mapped to the SPE token, which is inserted after BOS, giving the first layer *T* = *n* + 1 positions *t* = 0*, . . . , T* − 1,

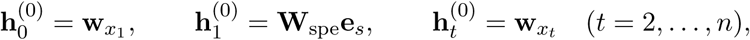

where 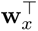 is row *x* of the token embedding matrix **W**_emb_ ∈ R^|V|×*d*^ and **W**_spe_ ∈ R*^d^*^×*d*s^ is learned.

### Decoder layers

Each layer comprises an attention sublayer and an MoE sublayer, each preceded by RMSNorm^35^ (pre-normalization) and wrapped in a residual connection. For *l* = 1*, . . . , L* and *t* = 0*, . . . , T* − 1,

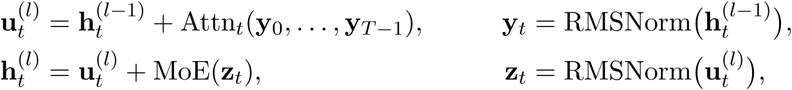

### Grouped-query attention

The *H* query heads form *G* groups of two consecutive heads, and each group shares one key head and one value head (grouped-query attention, GQA^31^). Causal scaled dot-product attention^69^ computes, for query heads *j* = 1*, . . . , H* and groups *m* = 1*, . . . , G*,

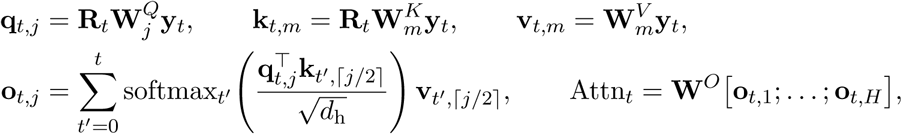

where 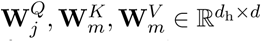 and 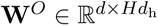 are the query, key, value and output projections, and softmax*_t′_* normalizes over *t*^′^ = 0*, . . . , t*, so each position attends only to itself and earlier positions. The matrix R_*t*_ ∈ R^*d*_h_^×^*d*_h_^ applies rotary position embedding (RoPE)^36^ at position *t* to queries and keys,

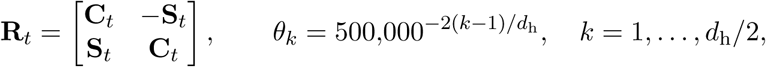

where **C***_t_*and **S***_t_*are diagonal with entries cos *tθ_k_* and sin *tθ_k_*, so coordinates *k* and *k* + *d*_h_*/*2 are rotated as a pair by the angle *tθ_k_*.

### Mixture-of-experts sublayer

Each MoE sublayer has one shared expert FFN^(s)^ and *E_l_* routed experts FFN^(r)^_*i*_ (i = 1, . . . ,*E*l), all SwiGLU feed-forward networks^48^ of the form FFN(**z**) = **W**_2_ (SiLU(**W**_1_**z**) ⊙ **W**_3_**z)** with expert-specific **W**_1_, 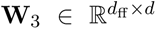 and 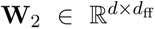 where SiLU(*ξ*) = *ξ σ*(*ξ*) acts elementwise and *σ* is the logistic sigmoid. The number of routed experts *E_l_* is 64, 32, 16, 8, 8, 8, 8, 8, 16, 32 and 64 for *l* = 2, 4*, . . . ,* 22 and 4 for odd *l*, 312 in total (Supplementary Fig. 3).

### Species-conditioned routing

The router W_r_ ∈ R^*E_l_*×*d*^ maps the MoE input **z** at a position to routing logits **W**_r_**z**. Feature-wise linear modulation (FiLM) ^38^ by the species embedding scales and shifts these logits before a softmax over all *E_l_* routed experts,

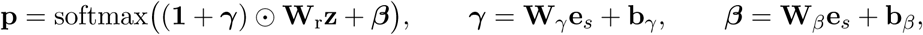

where **p** = (*p*_1_*, . . . , p_E_* )^⊤^ is the vector of routing probabilities, **W***_γ_,* 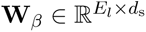 and **b***_γ_,* 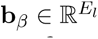 . As in the original FiLM implementation, the scale is parameterized as **1** + ***γ***, so that ***γ*** = ***β*** = **0** recovers the unmodulated logits. Because **e***_s_* is fixed for a sequence, ***γ*** and ***β*** are shared by all of its positions. At layer *l* and position *t*, **p** is written **p***_l_*(*t*), with entries *p_l,i_*(*t*) whose average over positions, *p̂_l,i_*, enters the auxiliary load-balancing loss.

Top-1 routing selects the routed expert *i*^∗^ = arg max*_i_ p_i_*, and only this routed expert is evaluated. As in Switch Transformer ^23^, its output is scaled by the gate value *p_i∗_* without renormalization. Pairing a shared expert, which processes every token, with one top-1 routed expert follows the residual MoE design^39^, with the two outputs combined by learned per-token mixing weights as in the DeepSpeed-MoE implementation,

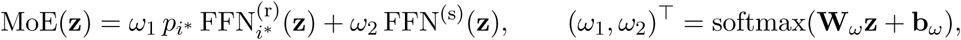

where **W***_ω_* ∈ R^2×^*^d^* and **b***_ω_* ∈ R^2^. Routing is dropless: no expert capacity limit is imposed, so every token is processed by its selected routed expert in training and inference, and the MoE sublayer acts on each position independently.

### Output

After layer *L*, a final RMSNorm is applied. The SPE position is removed from the returned hidden states, that is, this output and the layer inputs **h**^(0)^*, . . . ,* **h**^(^*^L^*^−1)^, so the SPE token never enters representation pooling. The output projection has no bias and shares its weights with the token embedding matrix (weight tying), so the next-token logits at position t ≠ 1 are W_emb_ RMSNorm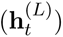. As the SPE output is removed and the BOS output is not scored, *x*_2_, the first token after BOS, is not a prediction target.

### MicroGlot-plain

MicroGlot-plain omits the SPE token and FiLM, that is, ***γ*** = ***β*** = **0** and **p** = softmax(**W**_r_**z**). Its input is 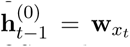 for *t* = 1*, . . . , n*; hence *T* = *n*, token *x_t_* occupies position *t* − 1 and the output at BOS predicts *x*_2_.

## Supplementary Note 4. Benchmark datasets

### 4.1 Deduplication

To reduce redundancy, we deduplicated each probing dataset separately by clustering at 90% nucleotide identity. The clustering tool was chosen according to sequence length. The viral family classification dataset and the marine, plant and synthetic species classification dataset, which contained sequences of up to 178 kb and 20 kb, respectively, were clustered with CD-HIT-EST^70,71^ (v.4.8.1) at 90% global nucleotide identity with a word length of 10. The geNomad, giant virus, BactoTraits and growth rate datasets contained chromosomes and genomes extending into the megabase range, which were too long for CD-HIT-EST. These datasets were therefore clustered with PSI-CD-HIT^72^, using megablast ^73^ from BLAST+^74^ (v.2.17.0) as the alignment program. The longest sequence in each cluster was retained as the cluster representative. For the geNomad and giant virus datasets, global sequence identity was used and all sequences were treated as linear. For the giant virus dataset, the contigs of each genome were concatenated into a single sequence, separated by 100 ambiguous bases (N). For the BactoTraits and growth rate datasets, individual contigs were clustered in fast mode, in which each contig is assigned to the first cluster that meets the threshold. Redundancy was then assessed at the genome level. A genome was considered redundant if contigs accounting for at least 90% of its total assembly length fell in clusters shared with another genome, and the shorter of the two genomes was removed. No additional deduplication was applied to the Genome Understanding Evaluation (GUE) datasets ^12^.

### 4.2 Replicon classification

We used the 625,420 geNomad reference sequences^44^, comprising 300,990 prokaryotic chromosome, 42,595 eukaryotic chromosome, 41,424 plasmid and 240,411 viral sequences. We excluded eukaryotic chromosome sequences, leaving 582,825 sequences across the three replicon classes. To balance the classes, we randomly subsampled each class to 41,424 sequences, the size of the smallest (plasmid) class. After deduplication, 121,769 sequences remained. These sequences were used for three tasks: viral versus non-viral classification (two classes); plasmid versus non-plasmid classification (two classes); and chromosome, plasmid and virus classification (three classes). The sequences were obtained from the Zenodo record accompanying geNomad^44^ (version 1.0, published 4 March 2023; https://doi.org/10.5281/zenodo.8049246).

### 4.3 Viral family classification

We used the known viral sequence cluster (kVSC) dataset from a human gut virome study ^46^, as distributed with Virgo^65^. Sequences assigned only at the class level were excluded, as were families with fewer than ten sequences before or after deduplication, leaving 473 sequences from eight families: *Aliceevansviridae*, *Autographiviridae*, *Drexlerviridae*, *Inoviridae*, *Madridviridae*, *Peduoviridae*, *Schitoviridae* and *Straboviridae*. The sequences were obtained from the figshare record of the Virgo benchmarking datasets^65^ (version 1, published 13 April 2025; https://doi.org/10.6084/m9.figshare.28730093.v1), which also contains the giant virus dataset.

### 4.4 Giant virus order classification

We used genome assemblies with order-level annotations from the Global Ocean Eukaryotic Viral (GOEV) database ^45^, as distributed with Virgo^65^. We retained the 1,412 genomes with available sequences, none of which was removed by deduplication. Of these, 1,370 genomes from four orders (*Algavirales*, *Asfuvirales*, *Chitovirales* and *Imitervirales*) were marked for inclusion in the source metadata and formed the dataset. Order labels were taken from Table_S4_GOEV_Genomic_database.xlsx, a supplementary table in the figshare record accompanying the GOEV database ^45^ (version 11, published 16 March 2023; https://doi.org/10.6084/m9.figshare.20284713.v11).

### 4.5 Marine, plant and synthetic species classification

We combined 12 datasets from the DNABERT-S benchmark ^13^, derived from simulated metagenomes of the second Critical Assessment of Metagenome Interpretation challenge (CAMI II)^66^ and from reference genomes. We merged species or operational taxonomic unit (OTU) labels shared across datasets, removed duplicate sequence–label pairs and randomly sampled up to 100 sequences per label, matching the per-species sampling limit of the source datasets. After deduplication, 110,492 of the 120,800 sequences remained, spanning 1,208 labels, of which 809 were OTUs. Of the 1,208 labels, 669 came only from the CAMI II marine datasets, 180 only from the CAMI II plant-associated datasets, 16 from both, and 343 from the synthetic datasets derived from reference genomes. The datasets were obtained from the DNABERT-S evaluation data archive^13^ (https://drive.google.com/file/d/1I44T2alXrtXPZrhkuca6QP3tFHxDW98c/view).

### 4.6 Phenotypic trait prediction

We matched species-level trait annotations from BactoTraits ^42^ to RefSeq genome assemblies ^25^, each comprising one or more contigs. BactoTraits encodes each trait by fuzzy coding, giving an affinity score for each of its categories (modalities). To obtain one classification label per species, we assigned each species to its highest-scoring category if that score, the mean of the strain-level affinities, was at least 0.5; ties between categories were resolved in favor of the category listed first in the BactoTraits file. Gram stain, motility and sporulation were treated as binary traits, whereas oxygen tolerance had four classes (aerobe, anaerobe, facultative and microaerophile), temperature preference had four (psychrophilic, psychrotrophic, mesophilic and thermophilic) and pH preference had five (acidophile, acido-neutrophile, neutrophile, alkalino-neutrophile and alkalino-alkaliphile). After label assignment, the matched datasets contained 9,071 genomes for Gram stain, 8,491 for motility, 4,096 for sporulation, 7,112 for oxygen tolerance, 14,208 for temperature preference and 2,063 for pH preference, together spanning 15,048 distinct genomes. To limit the size of the genome pool and prevent prohibitive computational cost, we sampled up to 300 genomes per class for each trait. Because temperature preference and pH preference each had classes smaller than this cap, every class of these two traits was further downsampled to the size of the smallest class, giving 202 and 123 trait annotations per class, respectively. The union of these per-trait samples comprised 3,777 genomes, of which 3,094 remained after deduplication. Each trait dataset then included all genomes in this pool with an annotation for that trait. The final datasets contained 2,499 genomes for Gram stain, 2,357 for motility, 1,351 for sporulation, 2,103 for oxygen tolerance, 2,899 for temperature preference and 890 for pH preference. The BactoTraits species- and strain-level annotation files are available at https://ordar.otelo.univ-lorraine.fr/files/ORDAR-182/BACTOTRAITS_DATASET_2026-01-28_SPECIESLVL.csv and https://ordar.otelo.univ-lorraine.fr/files/ORDAR-182/BACTOTRAITS_DATASET_2026-01-28.csv, respectively, in the OTELo ORDaR repository record https://doi.org/10.24396/ORDAR-182. Genome assemblies used for the phenotypic trait and growth rate tasks are available from NCBI Datasets under their versioned assembly accessions (https://www.ncbi.nlm.nih.gov/datasets/genome/).

### 4.7 Growth rate prediction

We obtained species-level minimum doubling times from the Phydon training data ^43^ and matched them to bacterial and archaeal genome assemblies with species assignments from GTDB release 220^37^. The regression target was the log_2_-transformed minimum doubling time in hours. After deduplication, 1,027 of the 1,473 available genomes remained. Doubling times are available from the Phydon bacterial training file ^43^ (dated 18 June 2024; https://github.com/xl0418/Phydon/blob/e3bd578a3eb4370b920e9779e5ad4278ead59c27/data-raw/GTDB_tax_trait_repGenome_in_tree_expanded.csv) and the archaeal training file (dated 4 July 2024; https://github.com/xl0418/Phydon/blob/3ebc16ba5bb0fad7327b039e87ccb685aaf30d4f/data-raw/GTDB_tax_trait_repGenome_in_tree_expanded_archaea.csv).

### 4.8 Taxonomic composition of the trait prediction datasets

After deduplication, we counted distinct named taxa from phylum to species in each trait prediction dataset, assigning each genome to one species. BactoTraits lineages were taken from the species-level annotation file, which uses the SILVA release 138.2 taxonomy^75^. For the growth rate dataset, we retained the species labels supplied by Phydon and resolved higher ranks by matching genome accessions to GTDB release 220. Subspecies were grouped with their parent species, and unresolved annotations were excluded. The GTDB release 220 bacterial and archaeal taxonomy files (released 24 April 2024) are available at https://data.gtdb.ecogenomic.org/releases/release220/220.0/.

### 4.9 GUE epigenetic marks prediction

The GUE benchmark from DNABERT-2^12^ includes ten binary classification datasets of 500-nucleotide yeast (*Saccharomyces cerevisiae*) sequences, one for each of H3, H3K4me1, H3K4me2, H3K4me3, H3K9ac, H3K14ac, H3K36me3, H3K79me3, H4 and H4ac (Supplementary Table 4). Each dataset is supplied with a random split into training, development and test sets containing 80%, 10% and 10% of the sequences, respectively. We retained these splits and used the development set for validation. All GUE datasets used in this study were obtained from the archive GUE_v2.zip^12^ (https://drive.google.com/file/d/1uOrwlf07qGQuruXqGXWMpPn8avBoW7T-/view), which contains both the GUE and GUE^+^ datasets. We followed the download instructions given by DNABERT-2 at https://github.com/MAGICS-LAB/DNABERT_2/blob/d87d29f5570f6d8238de734c932697ef657be59a/README.md.

### 4.10 GUE SARS-CoV-2 variant classification

The GUE SARS-CoV-2 dataset ^12^ comprises 999-nucleotide sequences from the GISAID EpiCoV database, each assigned to one of nine variants: Alpha, Beta, Gamma, Delta, Zeta, Eta, Iota, Kappa and Lambda. We retained the supplied training, development and test splits and used the development set for validation.

### 4.11 GUE viral and fungal species classification

The two GUE species classification datasets^12^ are derived from viral and fungal reference genomes in GenBank. The viral species dataset contains 5,000-nucleotide sequences from 25 species, and the fungal species dataset contains 10,000-nucleotide sequences from 20 species (Supplementary Table 4). We retained the supplied training, development and test splits for both datasets.

### Supplementary Note 5. Routing fingerprint and sequence composition

The routing analysis used 1,000 species from the pretraining corpus, 250 each from Bacteria, Archaea, Eukaryota and Viruses. Sequences were eligible if they were at least 8 kb long and carried no missing or placeholder labels (Supplementary Note 1) at any rank other than kingdom. Species with at least two eligible sequences were retained. Within each of the four groups, the 250 species were divided in balance among phyla, and the allocation of each taxon was divided in balance among its child taxa at each successive rank down to genus. When an allocation was smaller than the number of child taxa, the taxa receiving a species were chosen at random. When a taxon contained fewer eligible species than its allocation, the shortfall was reallocated among its sibling taxa. Species were selected at random within each genus. For each selected species, up to five of its distinct sequences were drawn at random, and the central 8,000-nucleotide window of each was used. All 1,000 species retained at least two windows, giving 3,948 windows in total. The selected species spanned three domains, five viral realms, 16 kingdoms, 85 phyla, 190 classes, 301 orders, 405 families and 615 genera.

Routing fingerprints were defined as follows. For window *w* and layer *l*,

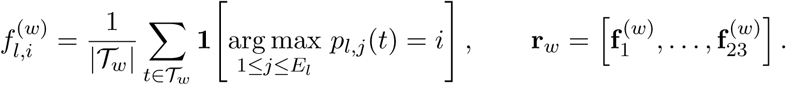

Here T*_w_* denotes the DNA content tokens of window *w*, defined as all tokens other than BOS, SPE and EOS. The quantity *p_l,j_*(*t*) is the routing probability of expert *j* at layer *l* for token *t*, that is, entry *j* of **p***_l_*(*t*) (Supplementary Note 3), and **1**[·] is the indicator function. Thus, 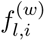 is the fraction of DNA content tokens whose top-1 routed expert at layer *l* is expert *i*, and the entries of each layer block 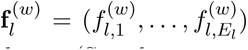 sum to one. The number of routed experts *E_l_* ranges from 4 to 64 across layers (Supplementary Fig. 3), so the concatenated routing fingerprint **r***_w_* has 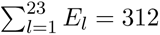 dimensions. Tokenized windows were limited to 4,096 tokens.

The flow diagram in Fig. 5a displays the 11 sparse layers with more than four routed experts. At each displayed layer *l*, window *w* was assigned to expert arg max_*i*_ 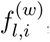, the expert selected most frequently among its DNA content tokens. Ribbons between consecutive displayed layers represent the windows assigned to each pair of experts, separated by the taxonomic group of the windows.

For the sequence composition comparison, tetranucleotide frequency (TNF) profiles were computed by counting overlapping tetranucleotides in each window, skipping those containing ambiguous bases, and merging each tetranucleotide with its reverse complement, following metagenomic binning ^51,52^. This gave 136 features, comprising 120 reverse-complement pairs and 16 tetranucleotides that are their own reverse complements, which were normalized to sum to one. Denoting either the routing fingerprint or the TNF profile of window *w* by **x***_w_*, we computed species-level feature vectors and standardized them as

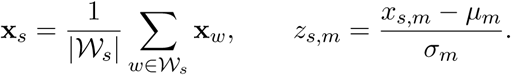

Here W*_s_* is the set of two to five windows from species *s*, with each window contributing equally, and *x_s,m_* is coordinate *m* of **x***_s_*. The quantities *µ_m_* and *σ_m_* are the mean and standard deviation of that coordinate across the 1,000 species, computed separately for each feature set. The standardized vectors **z***_s_* = (*z_s,_*_1_*, z_s,_*_2_*, . . .*) were used for UMAP visualization and *k*-means clustering in Fig. 5b, and those of the routing fingerprints were used for pairwise cosine similarities between species in Fig. 5c, where species were ordered by hierarchical clustering. For these analyses, Eukaryota was divided into fungi (97 species) and protists (153), and Viruses into the realms Riboviria (155), Duplodnaviria (44), Varidnaviria (41), Monodnaviria (9) and Adnaviria (1), giving nine reference groups together with Bacteria and Archaea.

## Supplementary Figures

**Supplementary Figure 1.**
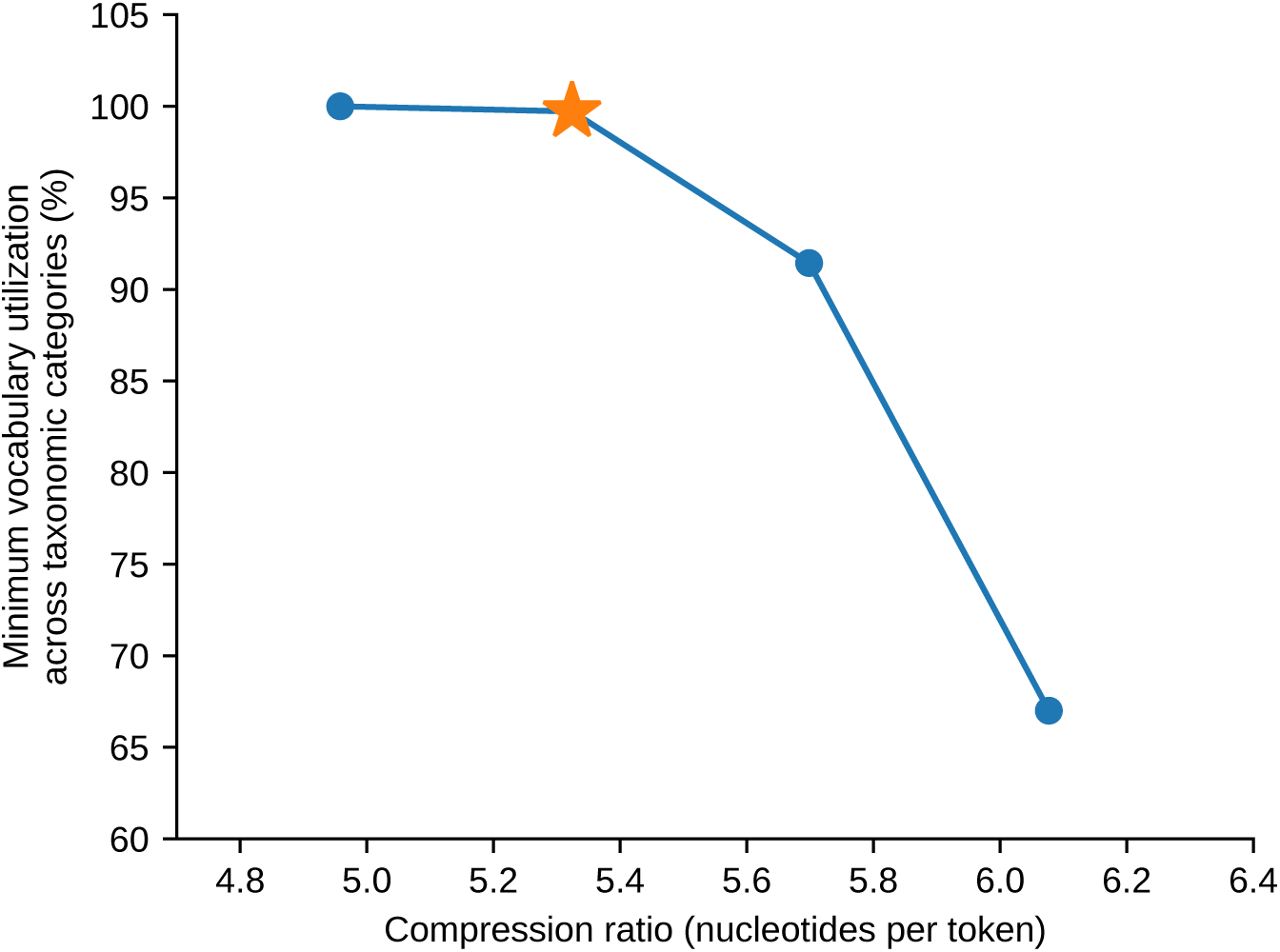
**Selection of BPE vocabulary size.** Compression ratio and vocabulary utilization for four candidate byte pair encoding (BPE) vocabularies of 4,096, 8,192, 16,384 and 32,768 tokens, shown from left to right. Compression is expressed as nucleotides per token. Vocabulary utilization is the percentage of vocabulary tokens observed at least 100 times within a taxonomic category. The minimum across six categories (bacteria, archaea, fungi, protists, viruses and plasmids) is shown. The star marks the selected vocabulary of 8,192 tokens.

**Supplementary Figure 2.**
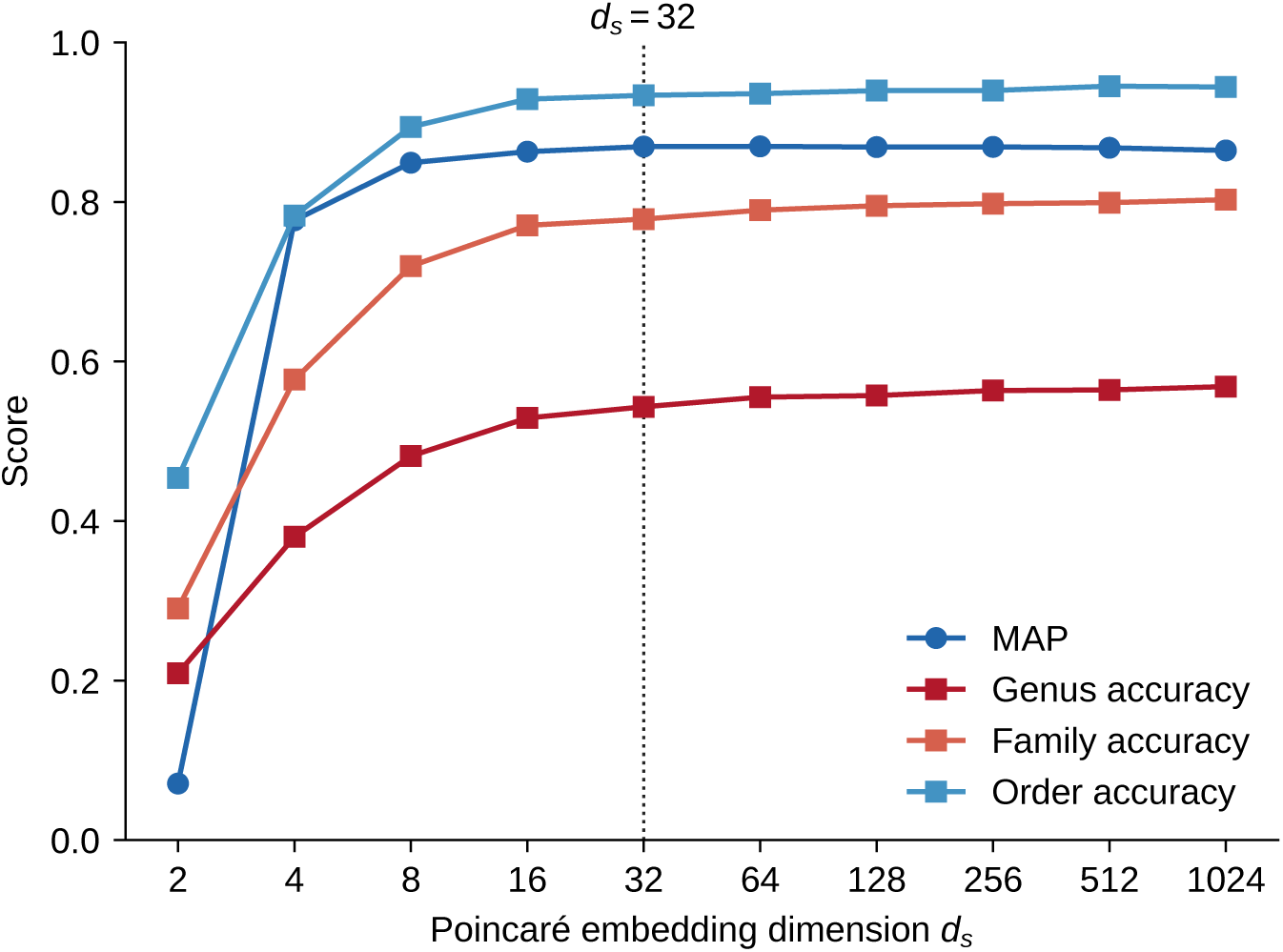
**Selection of Poincaré embedding dimensionality.** Reconstruction of taxonomic relationships and prediction of taxonomic ranks across embedding dimensions *d_s_*. Circles show mean average precision (MAP) for distance-ranked retrieval of taxonomic relationships ^30^, with higher values indicating better retrieval and MAP = 1 indicating perfect retrieval. Squares show linear probe accuracy at the order, family and genus ranks. The dotted vertical line marks the selected dimension *d_s_* = 32, beyond which improvements were limited.

**Supplementary Figure 3.**
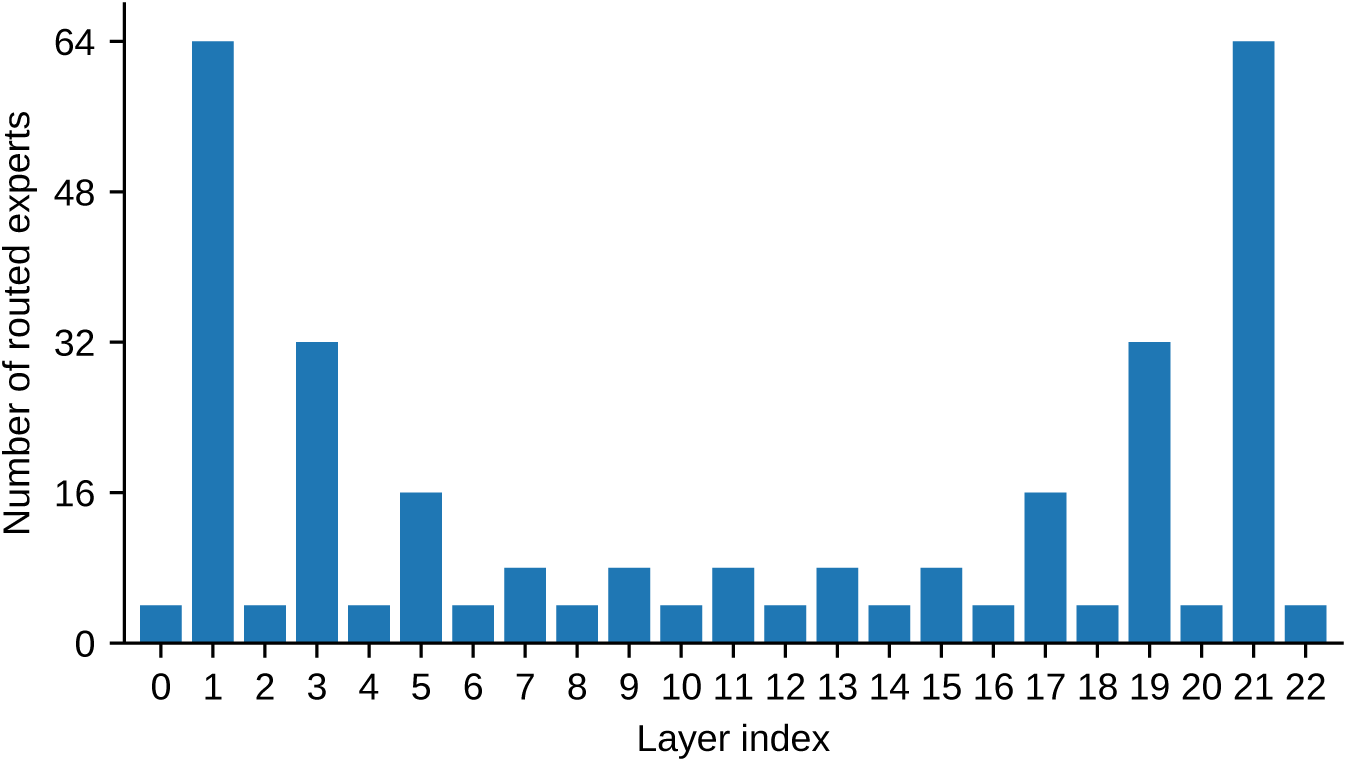
U-shaped allocation of routed experts across decoder layers. Number of routed experts in each of the 23 decoder layers of MicroGlot. Each layer also contains one shared expert, which is not shown.

**Supplementary Figure 4.**
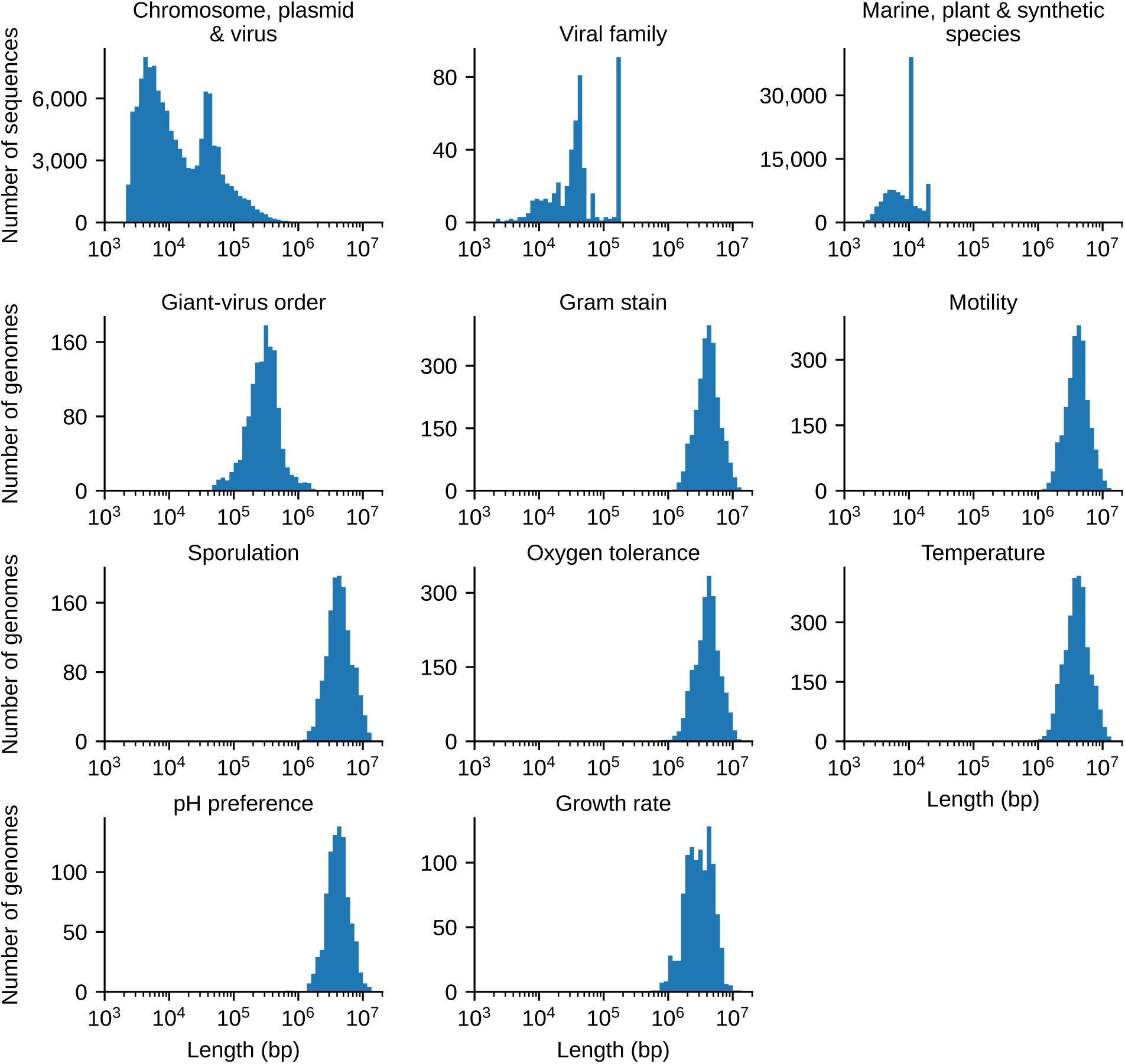
**Sequence length distributions of the probing datasets.** Histograms of sequence length for the datasets of the 13 probing tasks. The three replicon classification tasks share one geNomad dataset, shown in a single panel. The top row shows datasets of individual sequences. The remaining panels show datasets of genome assemblies, for which the length of each genome is its total assembly length. Lengths are shown on a logarithmic axis with logarithmically spaced bins.

**Supplementary Figure 5.**
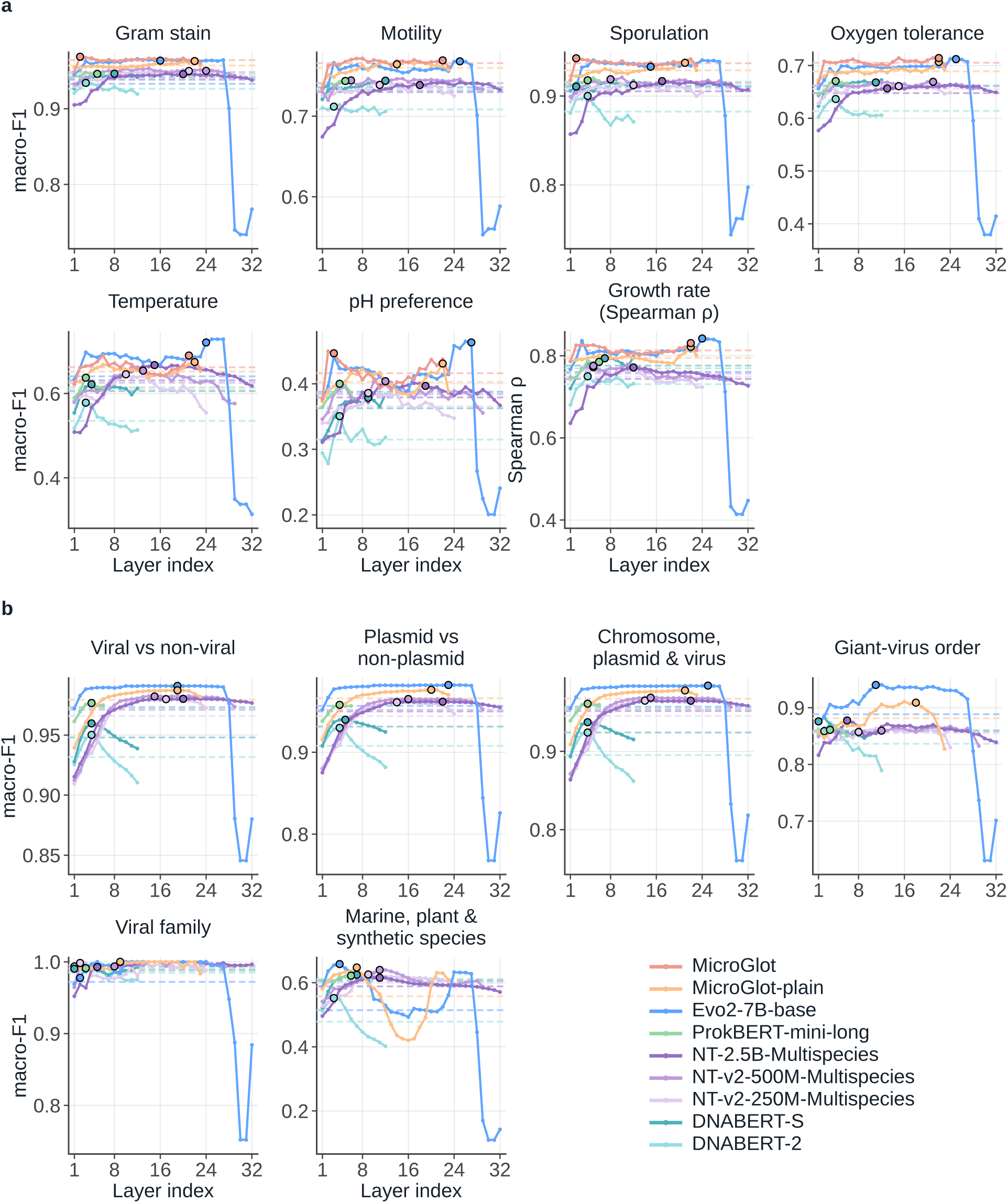
**Layer-wise probing of frozen gLM embeddings**. **a**, Seven trait prediction tasks, scored by macro F1 except growth rate, which uses Spearman *ρ*. **b**, Six microbial taxonomic classification tasks, scored by macro F1. Points show mean test scores across ten folds and three random seeds. Black outlines mark validation-selected best layers. Dashed lines indicate mean scores across layers.

**Supplementary Figure 6.**
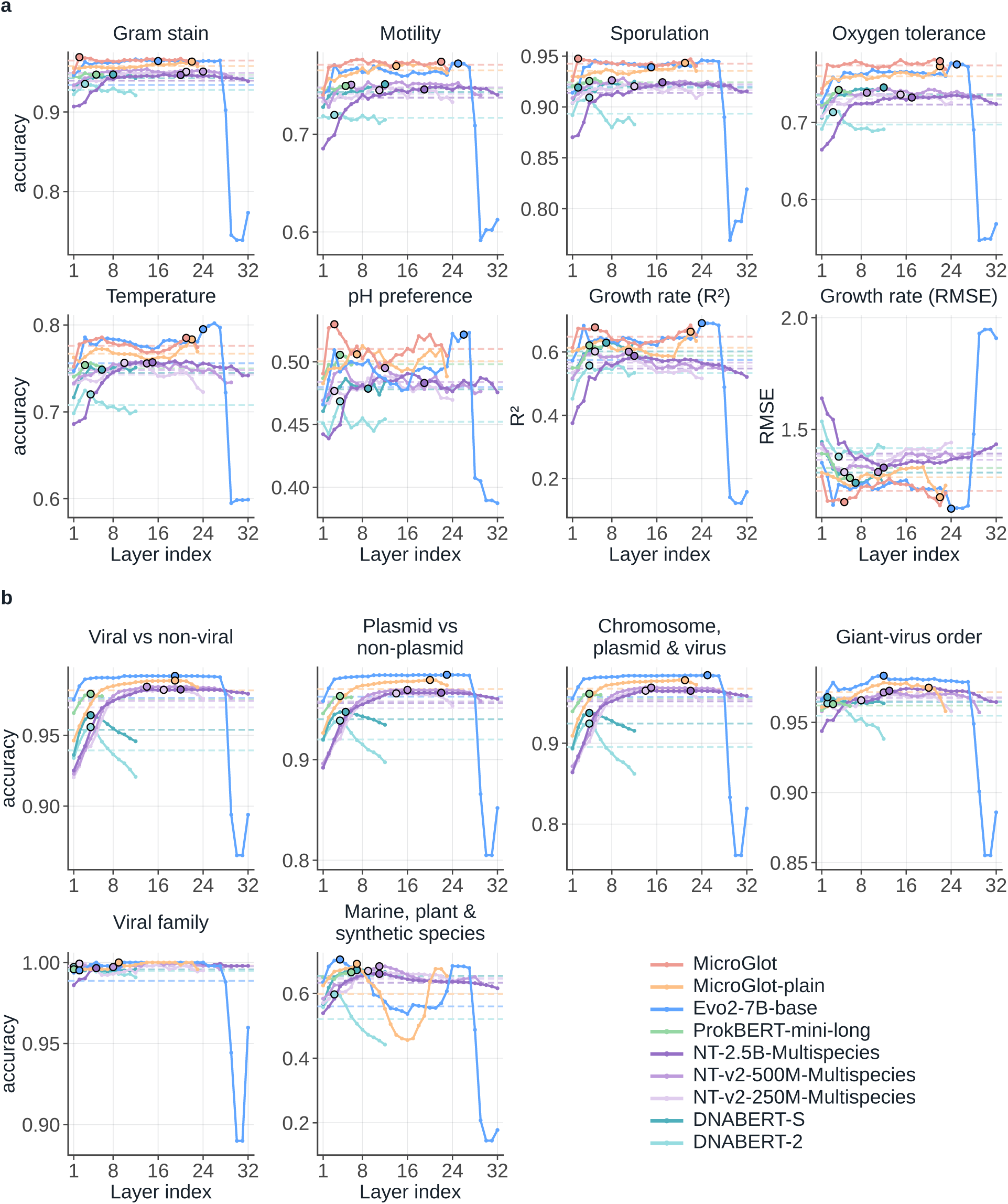
Layer-wise probing performance using secondary metrics. **a**, Accuracy for the six categorical trait tasks and the coefficient of determination (*R*^2^) and root mean squared error (RMSE) for growth rate prediction. **b**, Accuracy for the six microbial taxonomic classification tasks. Points show mean test scores across ten folds and three random seeds. Black outlines mark validation-selected best layers, chosen using the metric shown in each panel. Dashed lines indicate mean scores across layers. Lower RMSE indicates better performance.

**Supplementary Figure 7.**
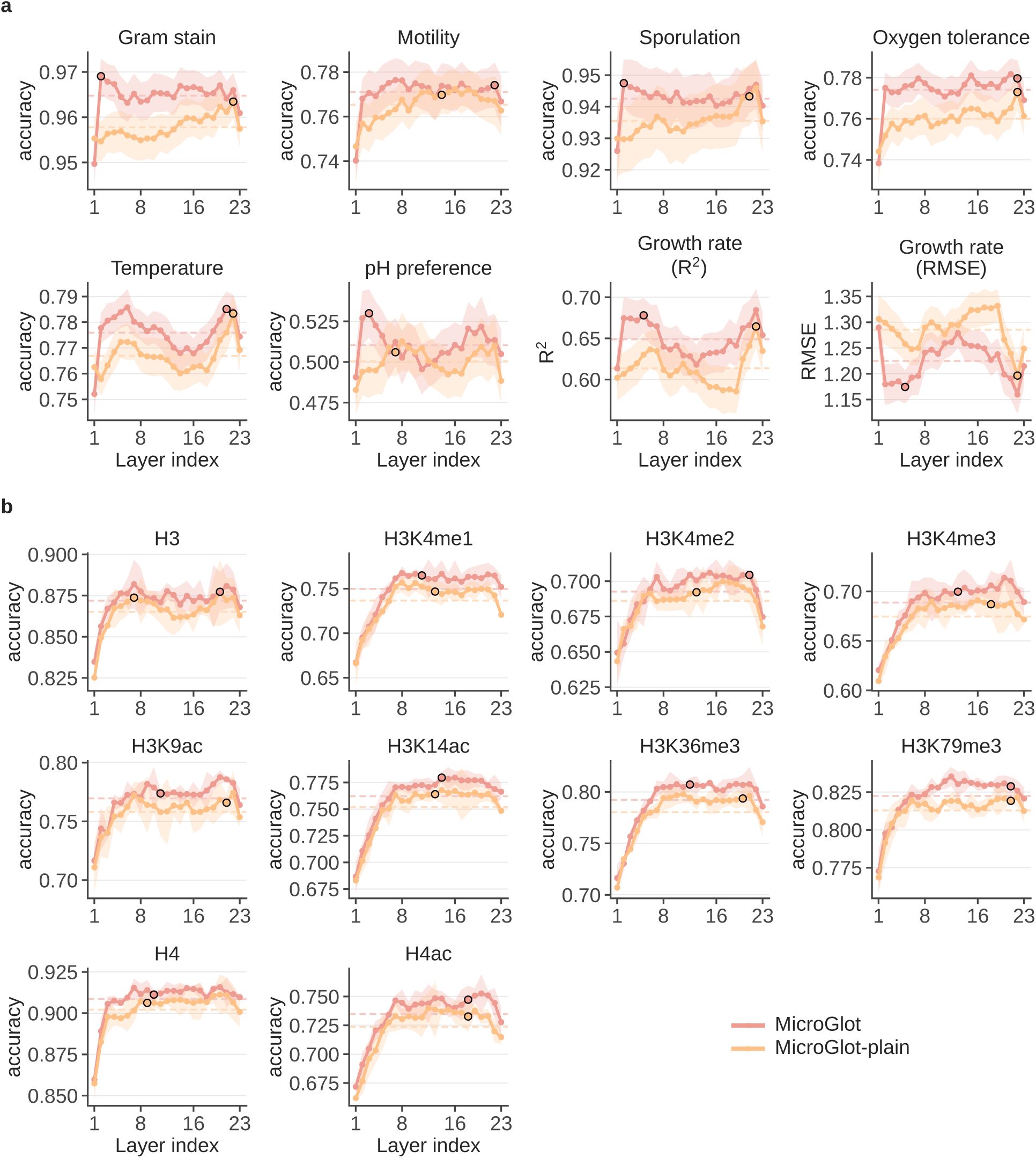
**Ablation of taxonomic information evaluated using secondary metrics.** Comparison of MicroGlot and MicroGlot-plain using the layer-wise probing experiments. **a**, Accuracy for the six categorical trait tasks and the coefficient of determination (*R*^2^) and root mean squared error (RMSE) for growth rate prediction. **b**, Accuracy for the ten epigenetic marks prediction tasks. Points show mean test scores at each layer, and shaded bands indicate 95% confidence intervals. Trait tasks used 10-fold cross-validation with three random seeds, and epigenetic marks tasks used the fixed benchmark split with three random seeds. Black outlines mark validation-selected best layers, chosen using the metric shown in each panel. Dashed lines indicate mean scores across all 23 layers. Lower RMSE indicates better performance.

## Supplementary Tables

**Supplementary Table 1.** MicroGlot architecture and model hyperparameters.

| Component | Specification |
| --- | --- |
| Backbone | Decoder-only Transformer |
| Attention | GQA (16 query heads, 8 key and value heads, head dimension 64) |
| Positional encoding | RoPE (base 500,000) |
| Normalization | RMSNorm (pre-norm) |
| Activation | SwiGLU (intermediate dimension 2,816) |
| Layers | 23 |
| Hidden dimension | 1,024 |
| Vocabulary | 8,192 (BPE) |
| Species embedding | 32 dimensions |
| Context | 8,192 tokens (∼44 kbp) |
| Total parameters | 2.98 B |
| Activated parameters | 479 M |
| Load balancing loss coefficient | 0.001 |

**Supplementary Table 2.** Configurations for MicroGlot pretraining.

| Hyperparameter | Value |
| --- | --- |
| Context length | 8,192 tokens |
| Optimizer steps | 22,087 (1 epoch) |
| Global batch size | ~4 M tokens |
| Optimizer | AdamW ($\beta_1=0.9, \beta_2=0.95$) |
| Peak learning rate | $3 \times 10^{-4}$ |
| Minimum learning rate | $3 \times 10^{-5}$ |
| Schedule | Cosine, 1% warmup |
| Weight decay | 0.1 |
| Gradient clipping (maximum norm) | 1.0 |
| Precision | bfloat16 (float32 routing and losses) |
| Parallelism | DeepSpeed ZeRO-1 |
| Gradient checkpointing | ✓ |

**Supplementary Table 3.**
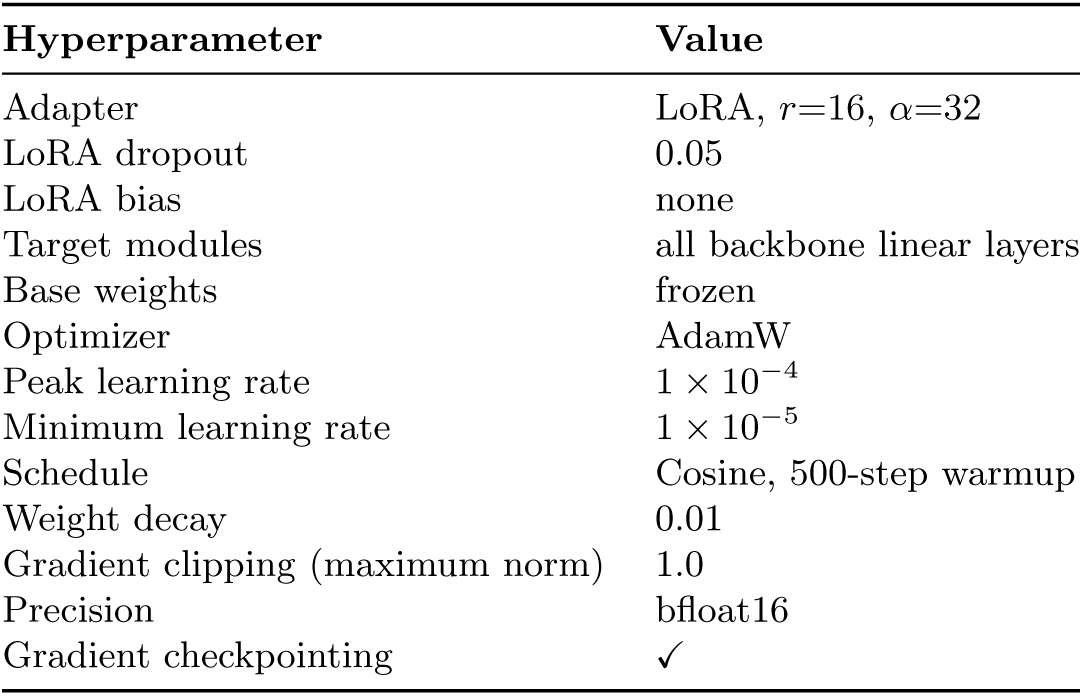
Configurations for LoRA fine-tuning.

**Supplementary Table 4.** Summary of GUE and GUE^+^ benchmark datasets for fine-tuning.

| Task | Classes | Length (nt) | Training | Development | Test |
| --- | --- | --- | --- | --- | --- |
| *Taxonomic classification* |  |  |  |  |  |
| SARS-CoV-2 variant | 9 | 999 | 73,335 | 9,166 | 9,168 |
| Viral species | 25 | 5,000 | 4,000 | 500 | 500 |
| Fungal species | 20 | 10,000 | 8,000 | 1,000 | 1,000 |
| *Epigenetic marks prediction* |  |  |  |  |  |
| H3 | 2 | 500 | 11,971 | 1,497 | 1,497 |
| H3K4me1 | 2 | 500 | 25,341 | 3,168 | 3,168 |
| H3K4me2 | 2 | 500 | 24,545 | 3,069 | 3,069 |
| H3K4me3 | 2 | 500 | 29,439 | 3,680 | 3,680 |
| H3K9ac | 2 | 500 | 22,224 | 2,779 | 2,779 |
| H3K14ac | 2 | 500 | 26,438 | 3,305 | 3,305 |
| H3K36me3 | 2 | 500 | 27,904 | 3,488 | 3,488 |
| H3K79me3 | 2 | 500 | 23,069 | 2,884 | 2,884 |
| H4 | 2 | 500 | 11,679 | 1,461 | 1,461 |
| H4ac | 2 | 500 | 27,275 | 3,410 | 3,410 |

